# Pooled screening for CAR function identifies novel IL13Rα2-targeted CARs for treatment of glioblastoma

**DOI:** 10.1101/2024.04.04.586240

**Authors:** Khloe S. Gordon, Caleb R. Perez, Andrea Garmilla, Maxine S.Y. Lam, Joey Jy Aw, Anisha Datta, Douglas A. Lauffenburger, Andrea Pavesi, Michael E. Birnbaum

## Abstract

Chimeric antigen receptor therapies have demonstrated potent efficacy in treating B cell malignancies, but have yet to meaningfully translate to solid tumors. Here, we utilize our pooled screening platform, CARPOOL, to expedite the discovery of CARs with anti-tumor functions necessary for solid tumor efficacy. We performed selections in primary human T cells expressing a library of 1.3×10^6^ 3^rd^ generation CARs targeting IL13Rα2, a cancer testis antigen commonly expressed in glioblastoma. Selections were performed for cytotoxicity, proliferation, memory formation, and persistence upon repeated antigen challenge. Each enriched CAR robustly produced the phenotype for which it was selected, and one enriched CAR triggered potent cytotoxicity and long-term proliferation upon *in vitro* tumor rechallenge. It also showed significantly improved persistence and comparable antigen-specific tumor control in a microphysiological human *in vitro* model and a xenograft model of human glioblastoma. Taken together, this work demonstrates the utility of extending CARPOOL to diseases beyond hematological malignancies and represents the largest exploration of signaling combinations in human primary cells to date.

## INTRODUCTION

Glioblastoma (GBM) is the most common form of primary malignant brain tumor and has a very poor prognosis, with current standard of care offering only marginal survival benefit ^1^. There are very few therapeutic options for this disease, owing in part to the tightly regulated blood brain barrier (BBB) impeding drug trafficking ^2–4^. T cells are one of the few cell types that are capable of trafficking across this barrier in order to carry out immune surveillance ^5,6^. This, along with T cells’ ability to establish persistent memory populations and to trigger epitope spreading against multiple antigens with heterogenous expression in coordination with the rest of the endogenous immune system ^7–16^, makes chimeric antigen receptor (CAR) T cell therapies a uniquely promising treatment strategy for glioblastoma. However, while CARs have been transformative in the treatment of relapsed and refractory acute B cell leukemia (B-ALL), large B cell lymphoma (LBCL), and multiple myeloma (MM), as evidenced by the FDA approval of 6 different CAR-T cell products targeting CD19 or BCMA for these indications, they have not yet shown evidence of long-term durable responses in the context of solid tumor diseases ^17–21^. This is largely due to the unique challenges posed by a solid tumor, including tumor antigen heterogeneity, barriers to physical trafficking to and infiltration into the tumor, and immunosuppression within the tumor microenvironment that impedes T cell persistence and anti-tumor function. Accounting for these factors in CAR design and discovery may inform better design principles and aid in translation of solid tumor targeted CARs.

CARs redirect native T cell killing machinery towards a target antigen of interest by linking extracellular recognition domains to intracellular immunostimulatory signaling domains that are most commonly derived from 4-1BB and/or CD28 and CD3ζ, as is the case for CD19- and BCMA-targeted therapies. For glioblastoma-targeted CARs, target antigens of interest that have been pursued in the clinic include EGFRvIII, HER2, IL13Rα2, B7-H3, CD147, GD2, and chlorotoxin, while others such as CSPG4 and CAIX have been reported in the literature ^22–30^. Additional targeting approaches have been shown to enhance function and overcome antigen heterogeneity, including implementing IF/OR gated logic and secreting bispecific antibodies (BiTEs) ^15,31^. Strikingly, ∼95% of CARs under consideration in the clinic, including those that target solid tumors, share the same signaling domains as those employed against hematological malignancies ^32^.

While effective for treating blood cancers, we posit that these signaling domains may induce suboptimal responses in the context of solid tumors, which may require different effector programs from those employed against B cell malignancies given their distinct immunological challenges. Hence, we propose that a more extensive exploration of signaling programs may unlock useful T cell functions and expedite the translation of CAR-T cell therapies to solid tumors. Until recently, large scale signaling explorations employing selections for function in mammalian cells have been difficult. Within the last few years, several groups, including our own ^33^, have pioneered pooled screening strategies in which immune cells are engineered to express a CAR and tagged with an untranslated DNA barcode, subjected to antigen exposure, and screened for a phenotype of interest, after which the barcode frequencies can be readout via next generation sequencing (NGS) ^34–38^. Collectively, we refer to this approach as CARPOOL. While other groups have employed this technique on the scale of hundreds to thousands of library members, these libraries are largely composed of a limited set of costimulatory domains in combination with CD3ζ, only representing a fraction of the comprehensive diversity of signaling inputs and combinations that could be useful in eliciting anti-tumor function.

Furthermore, while our initial demonstration of CARPOOL utilized Jurkat cells due to the ease with which they can be engineered and expanded, primary T cells offer a broader and more nuanced range of clinically relevant phenotypes available for use as selection criteria. Thus, we speculated that designing our screens to select for anti-tumor function in primary T cells may enable the discovery of effective solid tumor targeted CARs. To test this theory, we employed a 1.3×10^6^-member library of 3^rd^ generation CARs targeting IL13Rα2 and expressed in human primary T cells, from which we identified a novel CAR that elicits potent anti-tumor cytotoxicity and long-term persistence following selections for cytotoxic degranulation, persistence, and memory formation—all of which are expected to be critical features in targeting solid tumors. To our knowledge, this work represents the largest exploration of CAR signaling in human primary T cells described to date.

## RESULTS

### Library construction

A 2^nd^ generation 4-1BB based CAR targeting IL13Rα2 (which will be referred to as 13BBζ) has been reported to produce complete remission in patients with recurrent multifocal lesions that were heterogenous in antigen expression, with three experiencing transient complete remission ^11,39^. The target antigen IL13Rα2 is a high affinity IL-13 decoy receptor and cancer testis antigen that drives tumor invasiveness and is commonly overexpressed in glioblastomas in addition to a variety of other cancer types, including pancreatic cancer, breast cancer, and melanoma, while its endogenous expression is largely restricted to the testis ^40–45^. Given its demonstrated clinical efficacy, IL13Rα2 was selected as the target antigen against which we chose to employ our CARPOOL selection strategy.

To target IL13Rα2, we constructed a 3^rd^ generation CAR library that utilizes a tethered IL-13 cytokine with an E13Y mutation (to abrogate binding to IL4Rα, the co-receptor for the more commonly expressed IL-13 receptor, IL13Rα1) as the extracellular targeting moiety, as described by Brown et al ^11^. The CAR also included an IgG4 hinge with mutations in the Fc binding region (L235E, N297Q) and a CD4 transmembrane domain ^11,46^, after which the intracellular signaling domains were inserted. This was performed using Golden Gate assembly to shuffle 110 different signaling domains into three intracellular positions within the CAR at random, yielding a theoretical library size of 1.3×10^6^ unique CARs—each of which included an 18-nucleotide barcode in the 3’ untranslated region (UTR) followed by an IRES and EGFP sequence. To unbiasedly explore a large space of potential signaling inputs, the receptor signaling domains were derived from the same pool that was described previously ^33^, with the addition of viral immunoreceptor tyrosine-based activating motifs (ITAMs) and cytokine receptor signaling domains (**Supplemental Table 1**). Viral ITAMs have evolved to enhance the propagation of infected host cells and are capable of initiating activation, proliferation, and differentiation, all of which are known requirements for CAR signaling ^47^. Additionally, cytokine receptor signaling is one of three signals required for T cell activation and plays an important role in directing T cell differentiation, expansion, and persistence ^48^; indeed, one study showed that incorporating truncated IL2Rβ and STAT3 signaling could enhance proliferation, polyfunctionality, cytotoxicity, and reduce terminal differentiation ^49^. **Figure 1A** shows the architecture and composition of this library.

**Figure 1.**
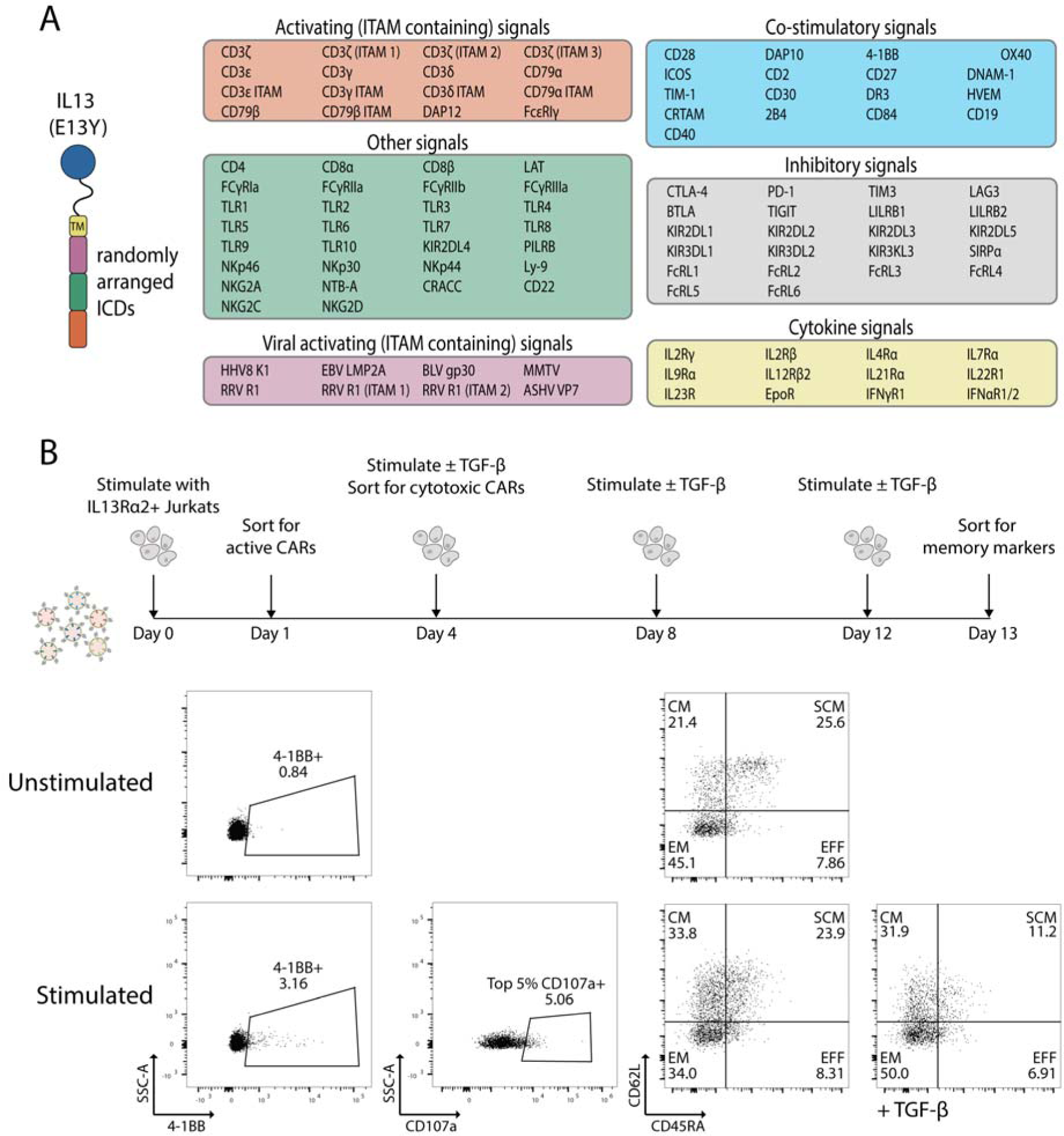
CARPOOL screening strategy. (**A**) Construct design for 1.3×10^6^-member library of IL13Rα2 targeted 3^rd^ generation CARs. (**B**) Rechallenge and sorting timeline for CAR enrichment. CARs were stimulated at each timepoint with growth arrested IL13Rα2^+^ Jurkat T cells at a 1:1 E:T ratio in 30 IU/ml IL-2 with or without 5 ng/ml TGF-*β* supplementation. Active CARs were sorted by collecting EGFP^+^ 4-1BB^+^ CAR-T cells, gated relative to unstimulated controls. Cytotoxic CARs were collected by sorting the top 5% of CD107a-expressing cells by MFI 6 hours after stimulation on day 4. Memory populations were sorted from stimulated CAR library on day 13 via CD62L and CD45RA expression. FACS plots are shown for sorted CD4^+^ library populations, with and without antigen stimulation. Data shown for donor 1.

While CD8^+^ T cells are known for their cytolytic functions, CD4^+^ T cells have been found to be more effective in treating glioblastoma *in vivo* and clinical evidence suggests that long-lived CD4^+^ memory T cells with sustained cytotoxic function have driven long-term remissions in patients treated with CD19-targeted CAR-T cell therapies ^50^. Thus, we decided to pursue selections in CD4^+^ human primary T cells. The resulting CAR-T cell libraries consisted of 1.0×10^7^ and 1.3×10^7^ CD4^+^ EGFP^+^ CAR-T cells, representing 7.7- and 10-fold library coverage, respectively. Notably, ∼26% of EGFP^+^ cells expressed CAR at the surface, as assessed by staining for the targeting ligand IL-13 (**Supplemental Figure 1A**).

### Library screening

To enrich persistent and proliferative CARs, CARPOOL transduced T cells were stimulated with growth-arrested IL13Rα2^+^ Jurkats every 4 days. Twenty-four hours after the initial antigen exposure, CAR-T cells exhibiting active responses were identified by the upregulation of the co-stimulatory molecule 4-1BB, which constituted 3.16% of the EGFP^+^ population, as demonstrated in **Figure 1B**. Cells displaying cytotoxic potential were enriched by sorting for the T cell degranulation marker CD107a following the second stimulation cycle (refer to **Figure 1B**). The remaining cells were rechallenged with or without TGF-*β* supplementation—which is commonly overexpressed in glioblastoma—to select for survival under immunosuppressive conditions ^51,52^. After the fourth round of antigen stimulation, T cells that sustained proliferative and persistent responses were collected. A subset of these cells was further segregated based on the expression of memory-associated markers CD62L and CD45RA, suggesting varying enrichment of memory T cell subsets (depicted in **Figure 1B**). This selection scheme—with the exception of CD107a sorting—was repeated with CD4^+^ T cells sourced from a second healthy donor (**Supplemental Figure 2**).

We used next-generation sequencing (NGS) to determine the frequency and identity of our selected CARs using previously described methods ^33^, where long-read sequencing was performed on day 13 rechallenged populations to maximize our ability to map ICD composition to barcodes in enriched pools. More than 50% of identified barcodes mapped to ICDs, with the majority containing 3 ICDs (**Supplemental Figure 1B**). Selected populations showed substantial enrichment in the most abundant barcodes relative to unselected population (**Figure 2A**), with the majority of enriched CARs containing ITAMs (**Figure 2B**). Co-stimulatory domains were also prevalent, while cytokine and inhibitory domains were relatively less enriched (**Figure 2B** and **Supplemental Figure 2C**). While 4-1BB and CD3ζ were among the top 30 ICDs for all selected populations, several others, including CD40, FCεRIγ, and DAP12, consistently enriched to a greater extent across biological replicates (**Figure 2C** and **Supplemental Figure 2D**). Notably, CD28 was not among the most frequent ICDs, which may be due to its proclivity to produce short-lived effector T cell responses ^53^. TGF-*β* treated stem cell memory and effector populations—which are grouped by CD45RA^-^ expression—shared many patterns of enrichment upon assessing the frequency of the top 30 ICDs, irrespective of position, in the selected populations (**Figure 2C**), but TGF-*β* supplementation had little effect on the relative abundance of particular signaling domains (**Figure 2C-D**).

**Figure 2.**
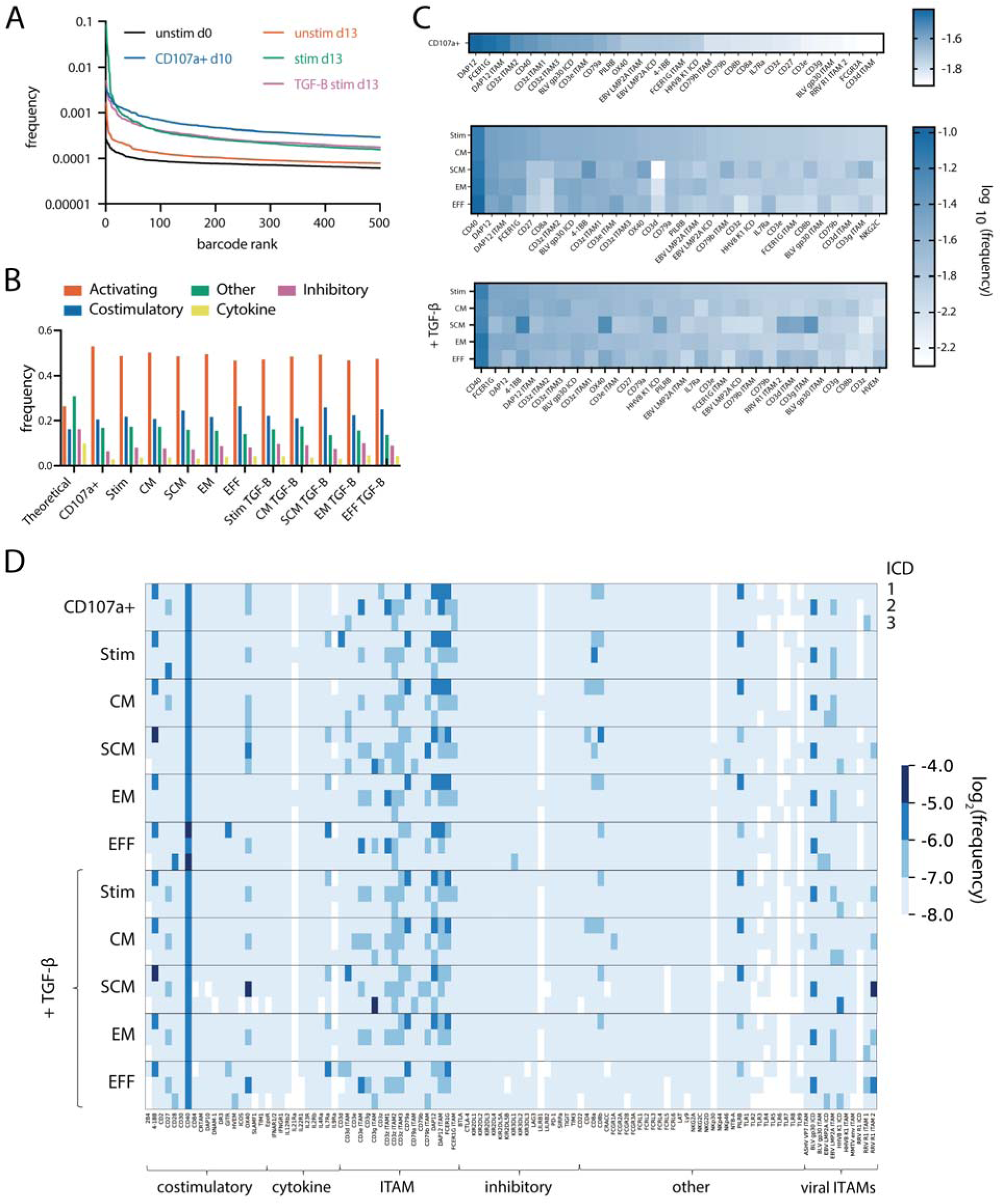
Sequencing analysis of enriched CD4^+^ library barcodes and signaling domains. **(A)** Barcode enrichment for unselected and selected populations, with barcode frequency on the y-axis and barcode rank (by frequency) on the x-axis. (**B**) Frequency of each family of signaling domain throughout different selected populations. (**C**) Heat maps showing bulk log_10_ frequencies of ICDs (irrespective of position) for CD107a sorted CARs and rechallenged CARs, with or without TGF-*β*. (**D**) Log_2_ frequencies of ICDs at each intracellular position relative to the transmembrane domain for selected populations. Data shown for donor 1. Barcode and ICD enrichment data can be found in Supplemental Data Files 1 and 2

During our examination of the spatial orientation of enriched ICDs, we discovered that most of them strongly prefer either one or two positions, with the exception of CD40. Additionally, we observed that 4-1BB is enriched in the membrane-proximal position, which aligns with the current design of CAR constructs that include 4-1BB. Other ICDs, some of which have not previously been tested in a CAR, consistently enriched in the same positions across CD4^+^ replicates, including CD79α, PILRβ, CD8α, and BLV gp30 (**Figure 2D** and **Supplemental Figure 2F**). While the top enriched ICD combinations differed between biological replicates, the patterns of ICD enrichment were largely reproducible between different batches of CAR library selections.

### CAR 4 exhibits potent anti-tumor function *in vitro*

Next, we selected 6 CARs that enriched in different selection criteria and harbored diverse ICD compositions (**Figure 3A**) to test for expression in Jurkat T cells, with 5 showing surface expression (**Supplemental Figure 3A**). We then tested the 5 selected CARs—which we named CARs 1, 2, 3, 4, and 5—along with a 13BBζ control for basal CAR expression, memory, and activation phenotype. We validated these constructs in both CD4^+^ and CD8^+^ human primary T cells given that both are used in the clinic and we hypothesized that these signaling perturbations may produce useful but distinct functions in CD8^+^ T cells. We found that 2, 4, and 5 showed slightly higher basal activity while CARs 2 and 5 produced a lower proportion of naïve cells, possibly due to tonic signaling (**Supplemental Figure 3C-E**). Upon expression in human primary CD4^+^ T cells and antigen challenge with IL13Rα2^+^ U87 cells—which were transduced to exogenously overexpress Flag-tagged IL13Rα2 (**Supplemental Figure 3F**)—at varying E:T ratios, CAR 4—which was the 8^th^ most enriched CAR in 4-1BB sorted cells, 1^st^ in CD107a sorted cells, 8^th^ in effector memory sorted cells, and 13^th^ in rechallenged cells—consistently showed robust activation via CD69 and 4-1BB upregulation as well as potent tumor cell killing in both CD4^+^ and CD8^+^ primary T cells (**Figure 3B-C**). Meanwhile, CARs 1, 2, and 5 showed lower activity, with CARs 2 and 5 being the only candidates to harbor ITIM-containing domains from the inhibitory proteins KIR3DL1 and TIM-3. While CAR 4 produced similar patterns of proinflammatory cytokine secretion to that of 13BBζ, including IFNγ and IL12p70 in both CD4^+^ and CD8^+^ T cells and Flt-3L and IL12-p40 in CD8^+^ T cells, CAR 3 was generally much less polyfunctional at an E:T ratio of 1:1 (**Supplemental Figure 4**). Interestingly, both CAR 3 and CAR 4 showed consistently lower levels of GM-CSF secretion in both CD4^+^ and CD8^+^ T cells and IL-1β in CD4^+^ T cells compared to 13BBζ (**Supplemental Figure 4**). At a higher E:T ratio of 1:10, CD4^+^ CAR 4 produced lower levels of GM-CSF in both CD4^+^ and CD8^+^ T cells, which is associated with CRS ^54,55^, along with similar levels of IFNγ, IL-17, and FLT3L in CD4^+^ T cells and similar levels of IFNγ in CD8^+^ T cells (**Supplemental Figure 4**).

**Figure 3.**
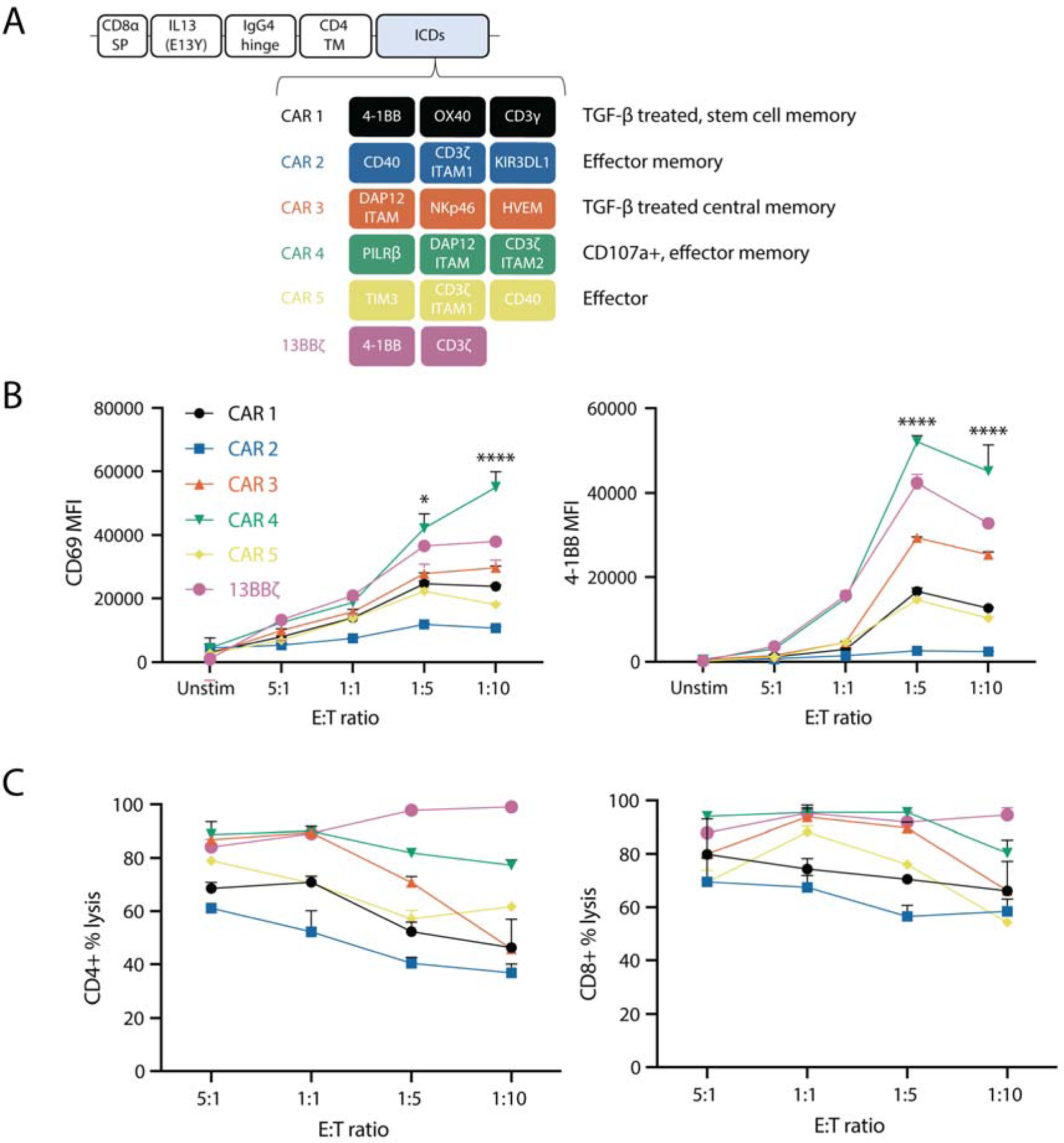
Enriched CARs show anti-tumor functions in response to glioblastoma *in vitro*. **(A)** Composition of enriched CARs selected for characterization, along with the populations in which they were enriched. Dose response curves following 24-hour co-culture of human primary CD4^+^ T cells with IL13Rα2^+^ U87 cells at varying E:T ratios (n = 3 technical replicates). Resulting (**B**) CD69 and 4-1BB upregulation in CD4^+^ T cells and (**C**) tumor cell killing in CD4^+^ and CD8^+^ T cells were measured after 24 hours. Data shown in (**B,C**) depicts means ± s.e.m. (n = 3 technical replicates). *P* values were determined using two-way ANOVA with Dunnett’s multiple comparisons test. For CD69 expression, *P* values were 0.0384 and <0.0001 for CAR 4 vs. 13BBζ at E:T ratios of 1:5 and 1:10 (n = 3, df = 60). For 4-1BB expression, both *P* values were <0.0001 for CAR 4 vs. 13BBζ at E:T ratios 1:5 and 1:10, respectively (n = 3 technical replicates, df = 59). Data in (**B**) is representative of two biological replicates, while data in (**C**) is representative of three biological replicates.

### Enriched CARs show enhanced proliferation relative to 13BB**ζ** upon rechallenge

In order to assess the long-term behavior of our selected CARs, we subjected them to four serial antigen challenges with IL13Rα2^+^ U87 cells on the same schedule described in **Figure 1B** with TGF-*β* supplementation and tracked CAR-T cell number at each time point, assessing memory and exhaustion phenotype on day 14. CAR 4 showed enhanced proliferation relative to 13BBζ by day 14 in two biological replicates, with no notable differences in memory phenotype (**Figure 4A-B** and **Supplemental Figure 5A**); however, CAR 4 showed higher levels of exhaustion marker expression (**Figure 4C** and **Supplemental Figure 5B**). Upon rechallenge with wild-type U87 cells expressing endogenous levels of IL13Rα2, we were intrigued to find that each CAR was enriched in the memory phenotype from which it was originally selected, with the exception of short-lived effectors (**Supplemental Figure 6**).

**Figure 4.**
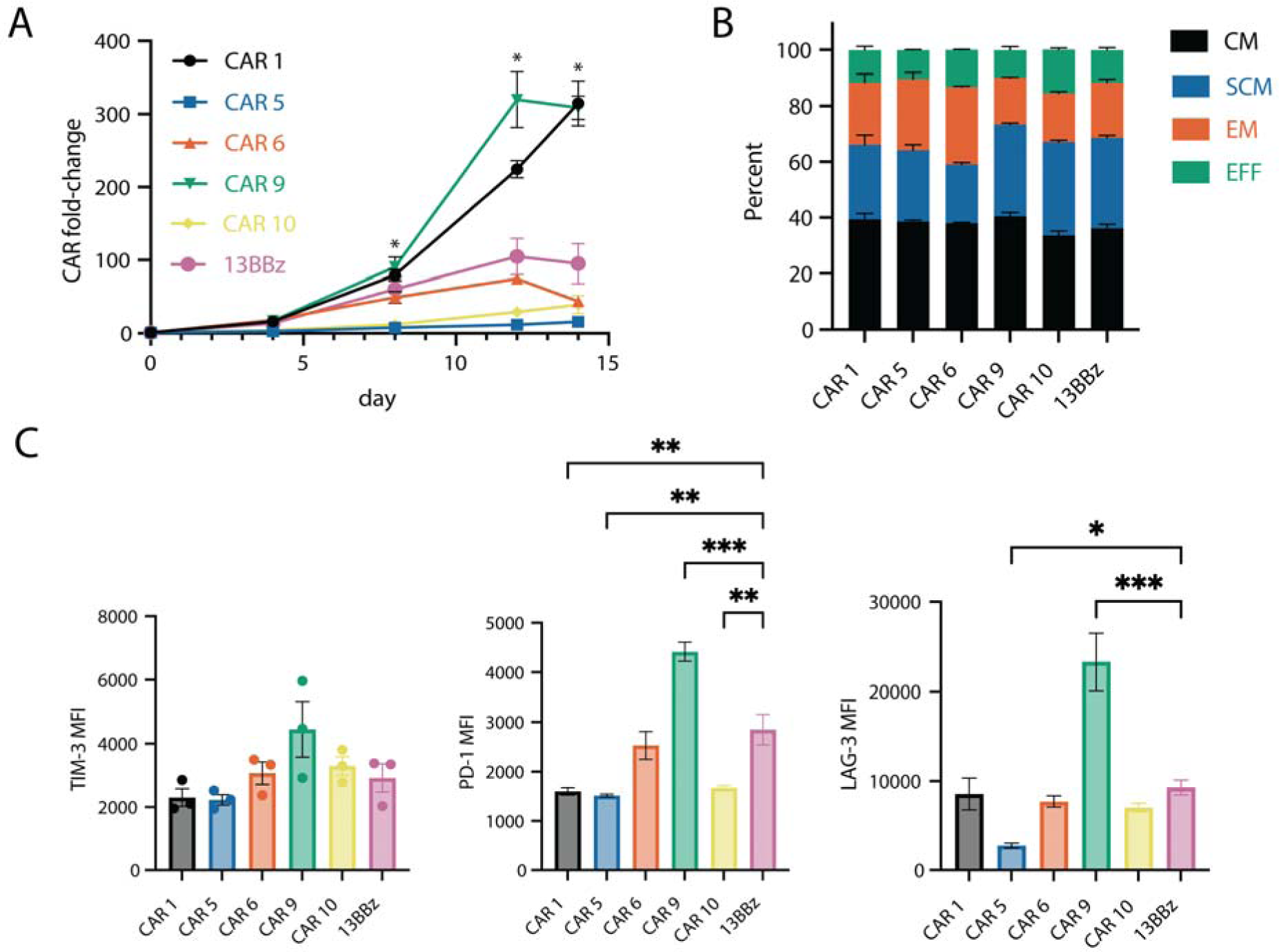
Selected CARs persist and proliferate in response to tumor rechallenge. (**A**) CAR fold-change following rechallenge with IL13Rα2^+^ U87 cells at a 1:1 E:T ratio on days 0, 4, 8, and 12. (**B**) Memory and (**C**) exhaustion phenotypes on day 14 of rechallenge. Data for (**A-C**) shows means ± s.e.m. (n = 3 technical replicates). The *P* values in (**A**) were determined using two-way ANOVA with Dunnett’s multiple comparisons test and are 0.0129, 0.0153, and 0.0141 for CAR 4 vs. 13BBζ on days 8, 12, and 14 (n = 3, df = 2). *P* values in (**C**) were determined using one-way ANOVA with Dunnett’s multiple comparisons test. In (**C**), the *P* values for PD-1 expression are 0.0024 for CAR 1 vs. 13BBζ, 0.0015 for CAR 2 vs. 13BBζ, 0.0004 for CAR 4 vs. 13BBζ, and 0.0038 for CAR 5 vs. 13BBζ. The *P* values for LAG-3 expression are 0.0466 for CAR 2 vs. 13BBζ and 0.0002 for CAR 4 vs. 13BBζ (n = 3 technical replicates, df = 12). Data is representative of two biological replicates.

### CAR 4 induces proliferative and cytotoxic transcriptional programs

Given that CAR 3 and CAR 4 showed the highest levels of activation and cytotoxicity, we next sought to determine whether their signaling inputs produced distinct transcriptional states. Thus, we subjected human primary donor CD3^+^ CAR-T cells to tumor rechallenge with IL13Rα2^+^ U87 cells, according to the timeline shown in **Figure 1B**. Then, we sorted the rechallenged cells for EGFP 48 hours after the final stimulation and performed single cell RNA sequencing. Dimensionality reduction and unsupervised clustering of the resulting transcriptional data yielded seven distinct cell clusters, with CAR 3 most enriched in clusters 1 and 7, CAR 4 enriched in clusters 2, 3, and 4, and 13BBζ enriched in clusters 4, 5, and 6—as confirmed by chi-squared analysis (**Figure 5A,B,D**). Notably, the CAR 4- and 13BBζ-enriched clusters 3 and 4 were enriched in dividing cells, which were largely CD8^+^ (**Figure 5C** and **Supplemental Figure 7**). Comparing average expression profiles, shown in **Figure 5E**, revealed that cluster 2, which was heavily enriched for CAR 4, contained genes related to cytotoxicity such as *PRF1*, *GZMK*, *GZMA*, *GNLY*, and *NKG7* along with *CCL5*—which distinguishes glioma infiltrating lymphocyte expansion within patient tumor samples when present in combination with a cytotoxic gene signature ^56,57^.

**Figure 5.**
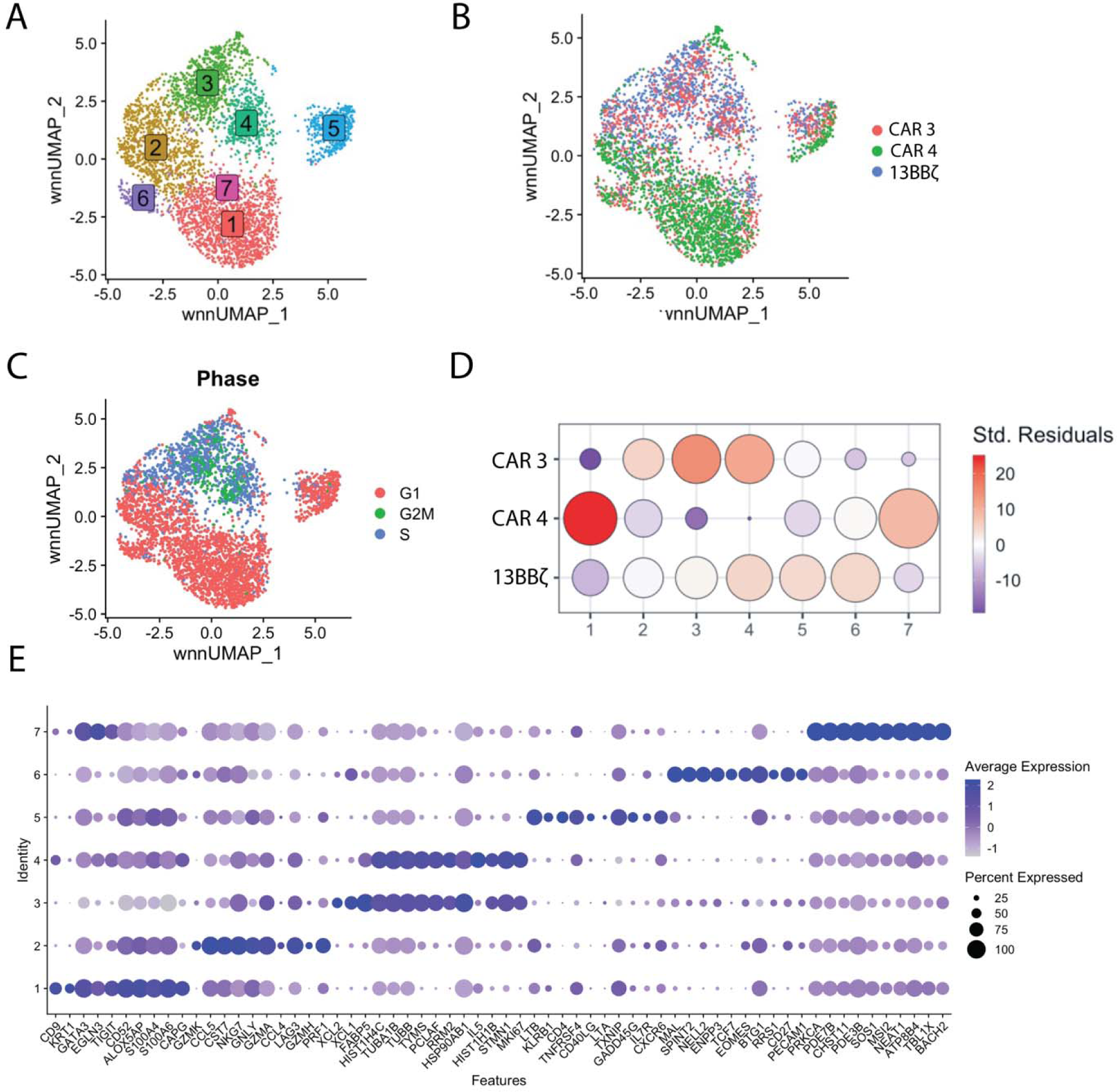
CAR 4 produces a cytotoxic gene signature that is associated with expansion within glioma patient samples. (**A**) UMAP embeddings of merged scRNA-seq profiles colored by cell state, (**B**) CAR identity, and (**C**) proliferative state following rechallenge of human primary CD3^+^ CAR-T cells with IL13Rα2^+^ U87 cells four times at an E:T ratio of 1:1 (n = 1471, 1471, and 1194 cells for CARs 3, 4, and 13BBζ, respectively). (**D**) Chi-square enrichment values for each CAR candidate within each cluster, represented by the Pearson residuals measuring the difference between the observed and expected CAR frequencies within each cluster. (**E**) Average expression of marker genes by cluster.

### CAR 4 elicits improved persistence and comparable tumor control *in vitro*

Given that CAR 4 exhibited both persistence and cytotoxicity, it became our lead candidate for subsequent characterization. We next sought to more accurately validate its anti-tumor activity compared to 13BBζ in a micro-physiological human *in vitro* model for GBM with a representative tumor microenvironment ^58^. Briefly, tumor spheroids were generated with IL13Rα2^+^ RFP^+^ U87 cells at ∼500 µm in diameter and seeded into a commercially available multi-well insert microfluidic device ^59^ in a fibrin gel with brain endothelial cells, pericytes, and astrocytes, and allowed to develop over 7 days as detailed in Lam et al. ^58^. The resultant model consists of a tumor spheroid with a dense core and invasive front, surrounded by a perfusable vasculature that mimics the *in vivo* BBB. The model would recapitulate physiological obstacles affecting CAR-T function, such as extravasation and tumor heterogeneity, providing a more accurate assay of CAR 4 function *in vitro*.

After formation of the GBM-BBB model, which we termed microtumors, 1×10^4^ CD3^+^ CAR 4 or 13BBζ CAR-T cells were perfused into the microfluidic devices. Tumor growth, CAR-T position, and proliferation were subsequently monitored over 9 days with confocal microscopy (**Figure 6A-D** and **Supplemental Figure 8**). Both CAR 4 and 13BBζ controlled tumor growth over 9 days and, while 13BBζ exhibited better tumor control on day 6, this regressed by day 9, at which point both CAR 4 and 13BBζ performed similarly in controlling tumor growth. (**Figure 6C-D** and **Supplemental Figure 8C**). Both CAR 4 and 13BBζ expanded in situ, as observed from an increase in signal over time, with CAR 4 expanding more by day 9 (**Figure 6D** and **Supplemental Figure 8B, D**). CAR 4 appeared to have better local tumor control at the invasive front, with a corresponding increase in CAR-T signal, suggesting local CAR-T function in controlling tumor growth. In contrast, 13BBζ tumor control did not correlate with locations of high CAR-T signal, suggesting more systemic control of tumor growth (**Figure 6C-D** and **Supplemental Figure 8C-D**). On day 9, T cells were retrieved from inside (IN) or outside (OUT) the microtumors (**Figure 6E**) and stained for T cell activation, exhaustion, and memory markers.

**Figure 6.**
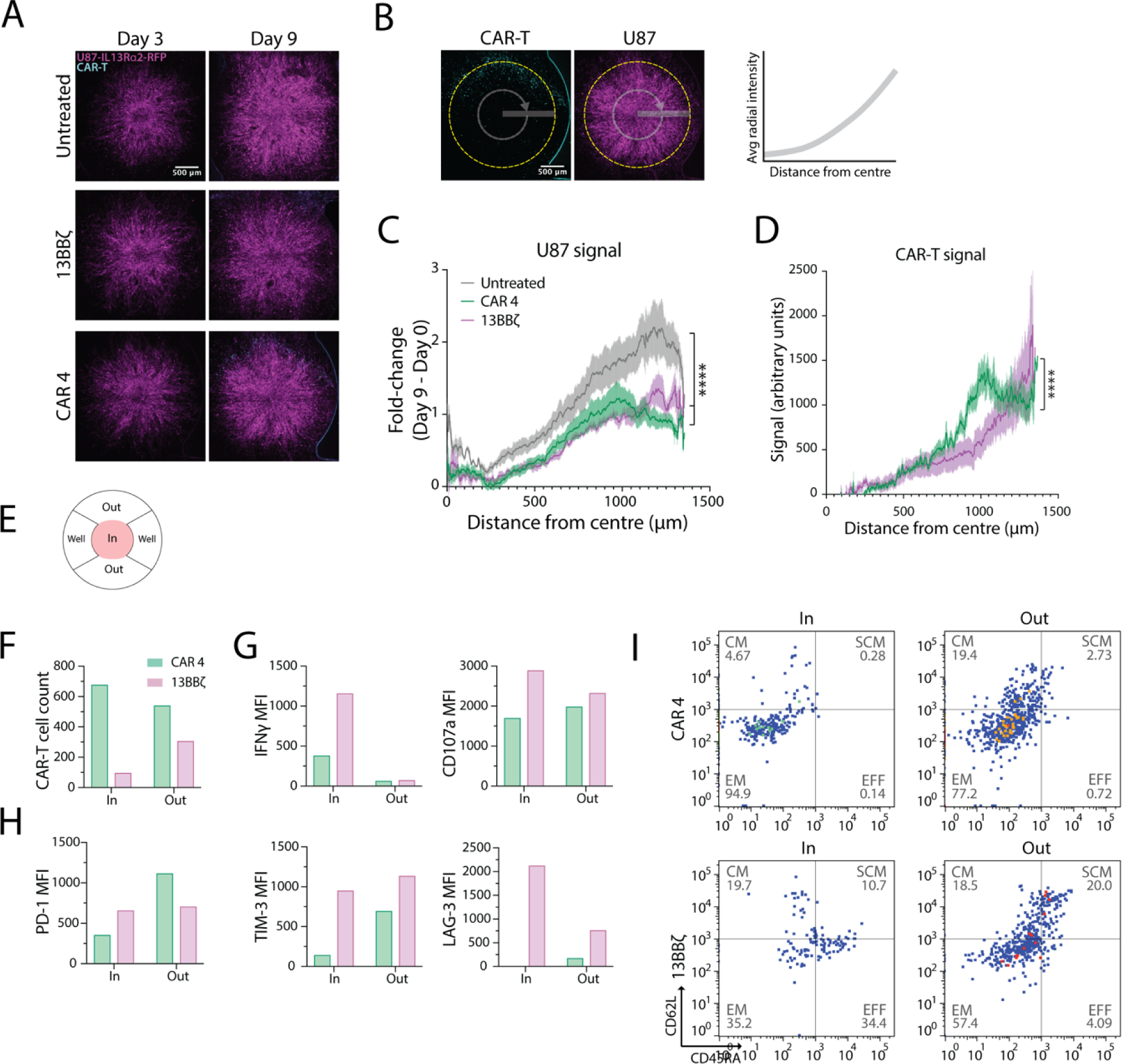
CAR 4 controls IL13Rα2^+^ U87 cells and exhibits increased persistence *in vitro*. **(A)** Representative image of IL13Rα2^+^ U87-RFP tumor spheroids (magenta) in microdevices at day 3 and day 9 after CAR-T (cyan) addition. (**B**) Illustration of radial plot analysis carried out in (**C**) and (**D**). (**C**) Increase in U87 cell signal intensity at day 9 relative to day 0. (**D**) CAR-T signal at day 9. Data shown in (**C**,**D**) are means ± s.e.m. (**E**) Illustration of regions inside and outside the microtumors. (**F**) CAR-T cell counts inside and outside the microtumors. (**G**) Activation, (**H**) exhaustion and (**I**) memory phenotypes of CAR-T cells retrieved from inside and outside the microtumors. *P* values in (**C**) were determined by comparing the area under the curve with one-way ANOVA with Tukey’s multiple comparisons test and are <0.0001 (n = 5 for untreated, 13BBζ and CAR 4, df = 12). *P* value in (**D**) were determined by comparing the area under the curve with two-way unpaired t-test and are <0.0001 (n = 5 for 13BBζ and CAR 4, df = 4). Data in (**F-I**) are pooled from 3 technical replicates.

Consistent with what we observed with live confocal microscopy, the T cell count for CAR 4 was higher than for 13BBζ, especially for T cells retrieved from inside the microtumors (**Figure 6F**). IFNγ expression was low outside the microtumors for both CAR 4 and 13BBζ, but increased after entering the microtumors, with 13BBζ having a bigger increase in expression (**Figure 6G**). CD107a expression was high for both CAR 4 and 13BBζ, with 13BBζ increasing and CAR 4 expression decreasing slightly upon entering the microtumors (**Figure 6G**). In contrast, exhaustion markers PD-1 and TIM-3 were highly expressed in both CAR 4 and 13BBζ outside of the microtumors, but had lower expressions inside the microtumors, possibly as a result of immunosuppression limiting upstream activation. Notably, this decrease was dramatic for CAR 4 but slight for 13BBζ (**Figure 6H**). On the other hand, LAG-3 expression was low outside of microtumors, and increased significantly for 13BBζ, but was almost absent in CAR 4 inside microtumors (**Figure 6H**). Both CAR 4 and 13BBζ expressed a mix of memory cells outside the microtumors, skewing towards effector memory and central memory subtypes, with 13BBζ having more effector and stem cell memory subtypes than CAR 4 (**Figure 6I**). Upon entering the microtumor, the majority of CAR 4 were effector memory populations, with almost no effector or stem cell memory subtypes. In contrast, 13BBζ lost their stem cell memory subtype but retained a mix of effector, effector memory and some central memory subtypes (**Figure 6I**). Taken together, this suggests that CAR 4 may be capable of persisting within the immunosuppressive tumor microenvironment while losing functionality relative to 13BBζ.

### CAR 4 elicits improved persistence and antigen-specific tumor control *in vivo*

Finally, we sought to assess the *in vivo* persistence and tumor control of our lead CAR candidate, CAR 4, relative to 13BBζ in a subcutaneous xenograft model of human glioblastoma. Mice were injected with 2×10^6^ Flag^+^ IL13Rα2^+^ FLuc^+^ U87 cells subcutaneously, followed by intravenous administration of 1×10^6^ or mock transduced T cells 21 days later (**Figure 7A**). Tumor area and flux along with weight change relative to pre-treatment as a proxy for toxicity were assessed for 20 days following ACT (**Figure 7B**, **Supplemental Figure 9A-B**, **Supplemental Figure 10**, **Supplemental Figure 11A-C**, and **Supplemental Figure 12**). On day 20 post ACT, CARs from tumors and spleens were harvested and stained for T cell memory and exhaustion phenotypes, along with antigen for tumor samples. This was performed using two different sources of donor T cells, designated as donors 3 and 4.

**Figure 7.**
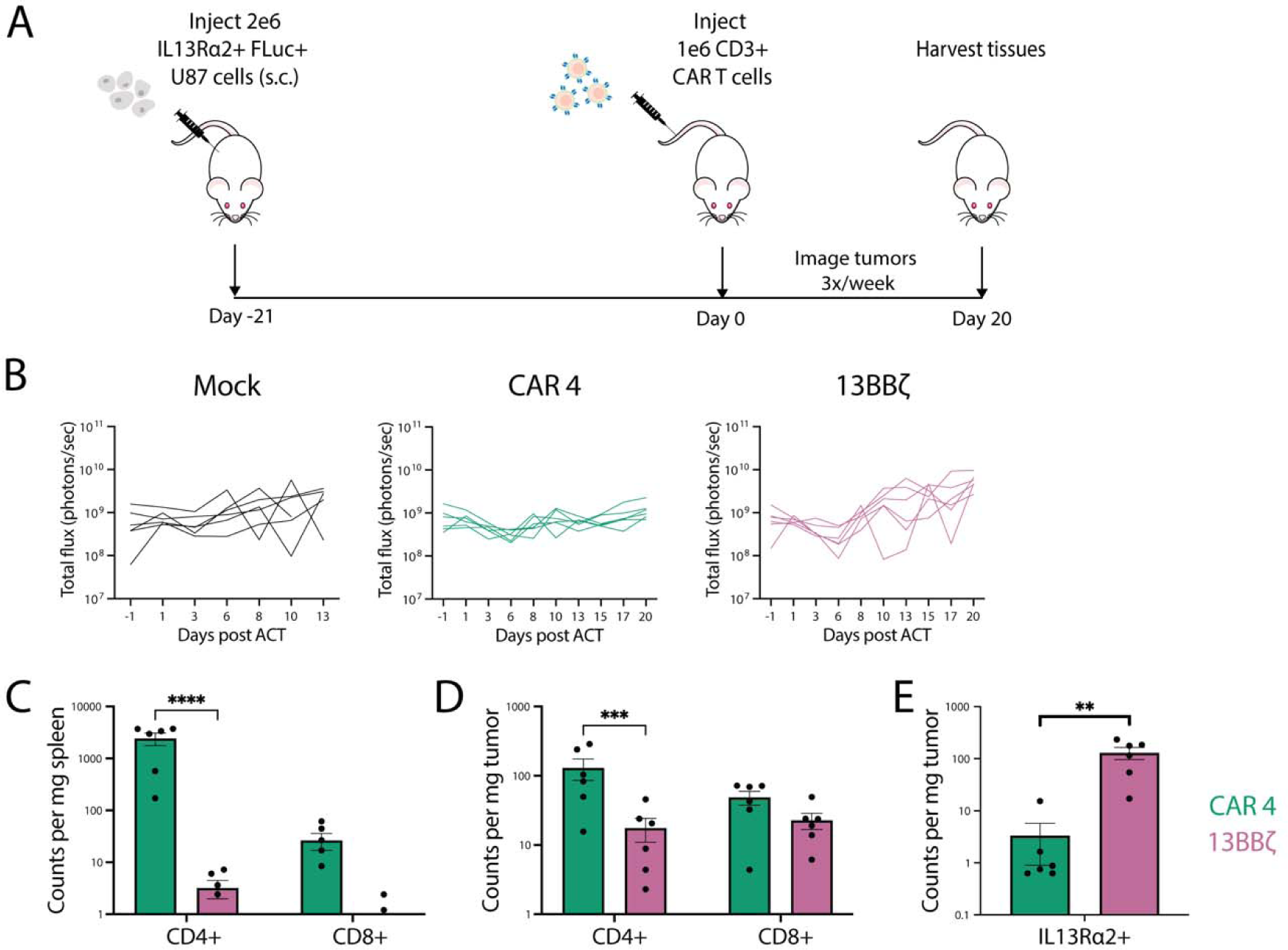
CAR 4 robustly controls IL13Rα2^+^ U87 cells *in vivo*. (**A**) Experimental timeline. (**B**) Tumor flux of Flag^+^ IL13Rα2^+^ FLuc^+^ U87 cells implanted subcutaneously in NSG mice. Data shown are for individual mice (n = 6 for mock, CAR 4, and 13BBζ). (**C**) CD4^+^ and CD8^+^ CAR abundance in the spleen and (**D**) tumor. (**E**) Antigen positive tumor cell abundances. Data shown in (**C-E**) are means ± s.e.m. The *P* values in (**B**) were determined using two-way ANOVA with Dunnett’s multiple comparisons test and are 0.0272, 0.0453, and <0.0001 for CAR 4 vs. 13BBζ on days 15, 17, and 20 respectively (n = 6 mice each, df = 180). *P* values in (**C**) and (**D**) were also determined by two-way ANOVA and are <0.0001 and 0.0003 for CD4^+^ CAR 4 vs. 13BBζ in the spleen and tumor, respectively (n= 6 each, df = 36). The *P* value in (**E**) is 0.0039 for CAR 4 vs. 13BBζ (n= 6 each, df = 5) as determined by an unpaired t-test. Data is representative of three biological replicates from donor 3.

Despite showing similar tumor control to 13BBζ, CAR 4 showed greater abundance in both the tumor and spleen for both donors (**Figure 7C-D** and **Supplemental Figure 11A,D-E**), alongside significantly lower IL13Rα2 expression within the tumor in donor 3 (**Figure 7E**); in donor 4-treated mice, most U87 cells following CAR treatment still retained low levels of antigen expression following CAR treatment relative to surviving mock-treated mice, suggesting potential antigen escape via downregulation (**Supplemental Figure 11F-G**). Notably, no CAR-treated mice experienced toxic side effects, as assessed by weight loss (**Supplemental Figure 9B** and **Supplemental Figure 11C**). In donor 3-treated mice, all CD8^+^ tumor-infiltrating CARs largely skewed towards effector phenotypes, while CD4^+^ CAR 4 produced a higher proportion of effector memory T cells relative to 13BBζ (**Supplemental Figure 9C,E**). In the spleen, CAR 4 showed greater persistence relative to 13BBζ for both donors, with a majority being CD4^+^ T cells that were overall less terminally differentiated than those found in the tumor (**Figure 7C** and **Supplemental Figure 9D,F** and **Supplemental Figure 11D,H-I**). In donor 4-treated mice, CAR 4 and 13BBζ produced similar proportions of memory populations in both niches (**Supplemental Figure 11H-I**). All tumor-infiltrating CARs showed similar levels of exhaustion marker co-expression, though CAR 4 showed significantly higher levels PD-1 in CD4^+^ and CD8^+^ T cells—which is thought to be an activation marker in the absence of LAG-3 and TIM-3—while 13BBζ showed higher expression of TIM-3 in CD8^+^ T cells in donor 3 (**Supplemental Figure 9G-H**). However, in donor 4-treated mice, CAR 4 showed comparable TIM-3 and LAG-3 expression (**S11J-K**).

## DISCUSSION

CARs have shown incredible efficacy in treating B-ALL, DLBCL, and MM, but have yet to translate to disease indications beyond hematological malignancies successfully. In this work, we utilized high throughput screening methods to select CD4^+^ CARs that persist, proliferate, differentiate, and kill tumor cells from a pool of signaling diversified CARs targeting the cancer testis antigen IL13Rα2. One CAR, consisting of PILRβ, DAP12 ITAM, and CD3ζ ITAM 2, was identified following selection for the cytotoxicity marker CD107a as well as persistence upon rechallenge and effector memory formation. It was found to produce significantly enhanced long-term proliferation and persistence upon tumor rechallenge compared to 13BBζ accompanied by elevated activation and cytotoxicity but lower secretion of GM-CSF—a cytokine implicated in producing CRS—*in vitro.* Given the high rate of recurrence in GBM patients, long-term persistence could prove to be pivotal in GBM targeted CAR-T cell therapies. It also showed a similar depletion of antigen-positive tumor cells in an IL13Rα2^+^ xenograft model of glioblastoma with markedly improved persistence in both the spleen and tumor—a feature which was also observed in a microphysiological human *in vitro* model of GBM with a tumor microenvironment. We also found that it robustly recapitulated effector memory formation in validation studies. This work demonstrates that our strategy is capable of identifying functional CARs from a large library that faithfully recapitulate the qualities for which they were selected.

Notably, the reduction in antigen load *in vivo* was somewhat subtle and did not correlate with a reduction in tumor size, likely due to the outgrowth of antigen-negative tumor cells; this phenomenon of downregulation is largely due to inherent variation in antigen expression within the tumor population where low antigen expression—possibly through differences in epigenetic state—can be selected for upon CAR-mediated tumor cell killing. In fact, the original case study using 13BBζ showed recurrence of antigen-negative tumor lesions, and several groups have employed dual targeting of both IL13Rα2 and EGFRvIII to overcome antigen heterogeneity and improve overall outcomes in treating glioblastoma ^16,31,60^; recent application of this dual targeting approach in phase I clinic trials has led to reduction in tumor burden ^61,62^. Our *in vitro* rechallenge studies employed mitomycin pre-treatment of the U87 cells in order to use them solely as a source of antigen and would not have captured this phenomenon, but future selections employing low antigen density target cells could increase CAR sensitivity in the context of antigen downregulation. The observed difference in translation between an *in vitro* and *in vivo* setting may be partly due to limitations in tumor infiltration as well as differential sensitivity to tumor immunosuppression, as seemed to be the case for CAR 4 upon assessment in a microfluidic device. Additionally, our immunocompromised xenograft model does not capture the interplay between the CAR-T cells and host immune system, a feature that is known to support CAR-T cell function as well as promote epitope spreading; the latter may be critical for targeting IL13Rα2 and treatment of glioblastoma in general.

CD3ζ ITAM 2 is commonly included in CARs via the full-length CD3ζ. While it has been examined in isolation ^63^, it has not been tested for function in a CAR at the membrane distal position of a 3^rd^ generation CAR. DAP12 is less commonly incorporated in the context of T cell-based therapies and its signaling is typically associated with NK and myeloid cells, where it plays an activating role in the context of high avidity ligands by phosphorylating LAT and NTAL and activating PI3K, PLCγ, VAV, ERK, and CBL signaling; it also initiates cytoskeletal reorganization and may enhance CD8^+^ T cell cytotoxicity ^64–72^. Meanwhile, PILRβ, an activator of SHP-1, has not—to our knowledge—been reported as functional in the context of a CAR and is typically expressed in NK cells where it causes activation in response to CD99 ligation; it is also thought to regulate T cell activation, but its exact role remains unclear ^73^. Further CAR domain deletion and protein-protein interaction studies may elucidate the exact mechanism of action and identify any synergistic function of this combination of signaling domains. While each of these domains would be plausibly useful in a CAR-T cell therapy upon examination in isolation, this exact combination and spatial configuration would be unlikely to emerge to the forefront through conventional design approaches.

While several groups have recently reported employing similar strategies for library-based screening of CARs ^34–36^, our study represents the most extensive exploration of signaling diversity in human primary T cells described to date in terms of signaling domain identity, functional family, immune cell source, and spatial arrangement; though screens of this scale in primary cells present unique technical challenges, their representative physiology enables use of a broader and more complex variety of screening criteria. While this large library size complicates comprehensive sampling of the represented molecular diversity, which would benefit the development of predictive machine learning algorithms and establishment of structure-function relationships for CAR design, it enables a more extensive search of the theoretical signaling landscape with fewer biases in inferring design principles. As such, we propose that this large library format may be a better tool for novel CAR discovery that can be broadly applied to many different disease targets and selection criteria. Altogether, the modular nature of high throughput CAR screening makes it uniquely poised to reinvigorate CAR discovery and expedite clinical translation in a streamlined process and its principles have broad implications for therapeutic modalities beyond CARs, making it a powerful tool for unlocking next generation cell therapies.

## METHODS

### Cell lines

HEK293T (CRL-3216) and Clone E6-1 Jurkat (TIB-152) lines were purchased from ATCC, while the U87 MG (HTB-14) line was generously gifted by Professor Forest White’s lab and transduced to stably express human IL13Rα2 with an N-terminal Flag tag along with firefly luciferase (FLuc) via an IRES sequence, then sorted for Flag and IL13Rα2 expression to produce the IL13Rα2^+^ U87 cell line (**Supplemental Figure 3F**). U87 MG IL13Rα2^+^ RFP^+^ cells were generated by stable expression of human IL13Rα2 and RFP (Addgene plasmid #26001) and sorting for RFP and IL13Rα2 expression. A target Jurkat T cell line was similarly produced by transduction to stably express IL13Rα2 and sorting for IL13Rα2 expression. HEKs and U87-derived cell lines were maintained in DMEM (ATCC) supplemented with 10% fetal bovine serum and 100 U/ml penicillin–streptomycin (Corning), while Jurkats were maintained in RPMI 1640 (ATCC) supplemented with 10% fetal bovine serum and 100 U/ml penicillin–streptomycin (Corning). Human brain microvascular endothelial cells were obtained from Angioproteomie (cAP-0002), maintained in EGM-2 MV BulletKit, and used at passage 6 (Lonza, CC-3156 & CC-4147). Primary human brain pericytes were obtained from Cell Systems (ACBRI 498), maintained in The System™ (Cell Systems, CSS-A101), and used at passage 6. Human astrocytes were obtained from ScienCell (1800), maintained in Astrocyte Medium from ScienCell (1801), and used at passage 4. Cell attachment factor (Cell Systems) was used to coat flasks for culturing of brain microvascular endothelial cells, pericytes, and astrocytes. Cell lines were routinely mycoplasma-tested using the MycoAlert PLUS Mycoplasma Detection Kit (Lonza).

### Plasmid construction

The plasmid pHIV-EGFP was gifted by Bryan Welm and Zena Werb (Addgene plasmid #21373), and pMD2.G and psPAX2 were gifted by Didier Trono (Addgene plasmid #12259 and #12260). To generate second-generation IL13Rα2 CAR-EGFP plasmid, a codon-optimized gene encoding IL13Rα2 CAR composed of E13Y mutated IL13 cytokine, L235E and N297Q-mutated IgG4 hinge, CD4 transmembrane domain, and ICDs derived from human 4-1BB and CD3ζ was PCR amplified from geneblocks purchased from IDT and cloned into the third-generation lentiviral vector pHIV-EGFP using Gibson Assembly. To generate a backbone vector for CAR plasmid library, the intracellular signaling domains of the IL13Rα2 CAR-EGFP plasmid were replaced with LacZ gene flanked by BsmBI restriction sites. The signaling-diversified CAR plasmid library was generated by PCR amplification of each ICD (**Supplemental Table 1**) at each of the three positions, with the forward and reverse primers adding unique linkers for each position. These products were then pooled at equimolar ratios for each position and combined with a pool of randomized 18mer barcode sequences. These were then inserted into a pGGA vector using BsaI restriction sites, with the insert also incorporating flanking BsmBI restriction sites. The final lentivirus plasmid library was generated by ligating the pGGA ICD library into the IL13Rα2 CAR backbone vector at the BsmBI restriction enzyme sites to replace the LacZ gene via Golden Gate Assembly. Final products were electroporated into DH10ß electrocompetent Escherichia coli cells (Thermo Scientific, EC0113) and purified to achieve a highly diverse plasmid library at ∼11.8x coverage of the theoretical diversity.

### Lentiviral production

Lentiviruses were generated by first transfecting 70% confluent HEKs with transfer plasmid, pMD2.g (VSVg), and psPAX2 combined at a plasmid mass ratio of 5.6:1:3 that was complexed with polyetherimide (PEI) at a DNA:PEI mass ratio of 1:3. For a confluent T225 flask, 42 μg of transfer plasmid was used for transfection. Medium was changed 3–6 hours after transfection, and lentiviral particles were collected in the supernatant 48–96 hours after transfection. The supernatant was then filtered through a 0.45-μm low-protein-binding filter, and centrifuged for 1.5 hours at 100,000 g. The pellet was then resuspended in serum-free OptiMEM overnight at 4°C and stored at −80 °C.

### Human T-cell activation, transduction, and expansion

Research using de-identified human blood was conducted as per MIT Committee on the Use of Humans as Experimental Subjects (COUHES) policies for exempt research. Peripheral blood mononuclear cells from healthy donors were purified from leukopaks purchased from Stem Cell Technologies using EasySep Direct Human PBMC Isolation Kits (Stem Cell Technologies) as per the manufacturer’s instructions. Primary CD4^+^, CD8^+^, or CD3^+^ T cells were isolated using EasySep Human CD4^+^, Human CD8^+^, or Human T Cell Enrichment Kits (Stem Cell Technologies) and cultured in RPMI 1640 (ATCC) supplemented with 10% fetal bovine serum, 100 U/ml penicillin–streptomycin (Corning), 30 IU/ml recombinant human IL-2 (R&D Systems), and 50 µM β-mercaptoethanol (Fisher). Before transduction, T cells were activated using a 1:1 ratio of DynaBeads Human T-Activator CD3/CD28 (Thermo Fisher) for 24 hours, after which 8 µg/ml of dextran (Sigma) and concentrated lentivirus were added to the culture at a multiplicity of infection of 5 for single lentiviral constructs and 2 for pooled library encoding lentivirus, where human primary CD4^+^ T cells were transduced at an efficiency of ∼10% to limit multiple integration events. Transduced cells were then sorted for EGFP expression. In the case of CAR library production, this was performed for two biological replicates using different donor PBMCs to source CD4^+^ T cells. After 3 days, DynaBeads and lentivirus were removed and cells were sorted for EGFP using a BD FACSAria II. Cells were rested for 4 days before characterization and maintained at a density of 5×10^5^ to 2×10^6^ cells/ml throughout.

### Flow cytometry and cell sorting

Cells were washed with 1× PBS (Sigma) supplemented with 0.5% bovine serum albumin (RPI) and 2mM EDTA, then surface stained by incubating with antibodies for 15 min on ice. They were subsequently washed again before flow analysis on a BD Accuri C6 or Beckman Cytoflex S or cell sorting with a BD FACSAria II or Sony MA900. Anti-IL13 (clone JES11-5A2), anti-CD213a2 (clone SHM38), anti-Flag (clone L5), anti-CD4 (clone SK3), anti-CD8 (clone SK1), anti-CD3 (clone OKT3), anti-CD107a (clone H4A3), anti-PD-1 (clone EH12.2H7), anti-TIM3 (clone F38-2E2), anti-LAG3 (clone 11C3C65), anti-CD62L (clone DREG-56), anti-CD45RA (clone HI100), and anti-CD69 (clone FN50) antibodies were purchased from Biolegend. For flow cytometry analysis of CAR-T cells isolated from microtumors, CAR-T cells in the side channels were flushed out of the devices and collected before microtumors were removed from microfluidic devices and incubated in a digestion mix consisting of 0.25% Trypsin, Nattokinase, collagenase IV, and DNAse for 30 min at 37°C, pipetting every 15 min. Liberated cells were then spun down and resuspended in 1× PBS (Sigma) supplemented with 0.5% bovine serum albumin (RPI) and 2mM EDTA, and stained accordingly. A BD FACSymphony™ A5 Cell Analyzer was used for acquisition. Anti-4-1BB (clone 4B4-1, Biolegend), anti-CD107a (clone H4A3, Biolegend), anti-IFN gamma (clone 4S.B3, eBioscience), anti-TNF-alpha (clone Mab11, BD), anti-CD62L (clone SK11, BD), anti-TIM-3 (clone 7D3, BD), anti-LAG-3 (polyclonal goat IgG, R&D Systems), anti-PD-1 (clone EH12.1, BD), and fixable viability dye eFluor 455UV (eBioscience) antibodies and dyes were used for staining. Data were exported and analyzed with FlowJo 10.10.0.

### CAR-T functional selections

In preparation for selections, 1.4×10^8^ human primary CD4^+^ T cells were transduced with lentivirus at a multiplicity of infection of 2 with 8 µg/ml dextran (Sigma). Virus and DynaBeads were removed after 3 days of transduction, and the cells were sorted for EGFP, with ∼10x library coverage, which was calculated based on the theoretical maximum diversity from the previous round, being maintained throughout. For a round of selection, cells were stimulated with mitomycin-treated IL13Rα2^+^ Jurkat cells, following 1 hour incubation in 50 µg/ml mitomycin at 37°C, then stained for 4-1BB after 24 hours or CD107a expression after 6 hours. For CD107a staining, cells were stimulated in brefeldin (BD 555029), monensin (BD 554724), and anti-CD107a antibody (clone H4A3) per the manufacturer’s instructions. The 4-1BB^+^ cells, gated relative to unstimulated library as a negative control, were collected after the first round of stimulation. After the second round, the top 5% of CD107a expressing T cells by mean fluorescent intensity were sorted on a BD FACS Aria II. Cells were then rested without antigen and expanded for 4 days before subsequent rounds of selection. After the final round of stimulation, 30-50% of cells were stained for CD62L and CD45RA and sorted for memory populations while the remaining unsorted cells were frozen for NGS. NGS sequencing data was deconvoluted and analyzed using a custom package called DomainSeq, as described in the manuscript.

### In vitro rechallenge assay

1×10^6^ CAR-transduced CD4^+^ T cells were co-cultured with mitomycin-treated target U87 cells, prepared by incubation in 50 µg/ml mitomycin for 1 hour at 37°C, at an effector to target (E:T) ratio of 1:1 in IL-2 deficient medium in triplicate. Every 2–3 days, approximately 5% of the culture volume was taken out for flow analysis for EGFP following addition of CountBright Plus Absolute Counting Beads (Thermo Fisher C36995). Then, 1×10^6^ CAR-T cells were taken out from the original culture and re-plated with a fresh batch of mitomycin-treated U87 cells at a 1:1 E:T ratio. CAR-T cells were sampled for scRNA-seq analysis at day 14, which was 48 hours following the fourth U87 challenge. On day 13, cells were also stained for memory markers (CD62L and CD45RA) and exhaustion markers (PD-1, TIM3, and LAG3).

### Cytotoxicity assay

U87 cells expressing FLuc were co-cultured with CD4^+^ or CD8^+^ T cells for 24 hours in IL-2-deficient medium at various E:T ratios. Cells were then collected and washed before cell lysis and addition of luciferin substrate from the Bright-Glo Luciferase Assay System (Promega). The resulting luminescent signal was measured using a Tecan Infinite M200 Pro. Signals were normalized to negative controls containing only target cells.

### Cytokine secretion assay

Following 24 hours of stimulation of human primary CAR-T cells with U87 cells, polyfunctional cytokine and chemokine secretion profiles in response to tumor challenge were determined using the 41-plex MILLIPLEX MAP Human Cytokine/Chemokine Magnetic Bead Panel from Miltenyi and measured on a Luminex FlexMap 3D system.

### PacBio and Illumina sequencing

Genomic DNA from selected cells was purified using the PureLink Genomic DNA Mini Kit (Thermo Fisher). For PacBio sequencing, PCR amplicons encoding CAR signaling domains and barcode regions were attached with SMRTbell adaptors using the SMRTbell Template Prep Kit 1.0 (Pacific Biosciences) and sequenced using a PacBio Sequel system. For Illumina sequencing, barcode regions were PCR amplified to conjugate P5 and P7 adaptor sequences and sequenced on an Illumina NextSeq500 system.

### Single-cell sequencing

Primary human T cells expressing CARs 3, 4, and 13BBz were sampled from U87 co-cultures following four tumor re-challenges, as described above. Samples were separately stained with a panel of CITE-seq antibodies (Biolegend) specific for a panel of T cell phenotypic markers, described previously ^74^, along with three unique hashing antibodies to label each CAR sample (Biolegend). Following staining and washing according to manufacturer’s protocols, live CAR^+^ cells were enriched from each sample on the basis of viability dye staining and GFP expression. Sorted samples were then pooled at approximately equal proportions, then encapsulated in two channels via Chromium Next GEM Single Cell 5’ Kit v2. Gene expression (GEX) and feature barcoding (FB; for CITE-seq antibodies) libraries were constructed based on manufacturer’s instructions, then pooled to achieve approximately 12.5% FB and 87.5% GEX, before sequencing on an Illumina NextSeq500 to a depth of 34,380 reads per cell. Reads were aligned to the Genome Reference Consortium Human Build 38 (GRCh38), and a cell-gene matrix was generated from the aligned reads using the CellRanger software (10X Genomics; v7.0.1). Analysis of this matrix was performed with the Seurat package (v4.3.0.1; in R v4.2.2) ^75,76^. To filter out low-quality or dying cells, cells with >5% of reads relating to mitochondrial genes were removed, as well as any cells with less than 1000 unique genes detected. Filtered cells were then assigned sample identity using hashtag antibody reads via the HTODemux algorithm ^76^; only cells confidently assigned to a single sample were used in downstream analysis. Although we aimed to encapsulate approximately equal numbers of cells from each sample, the data showed unequal cell calls; to address this, we downsampled cells to gain approximately equal cell depth per CAR, yielding 1471, 1471, and 1194 cells expressing CAR3, 4, and 13BBz, respectively. This downsampled dataset was normalized and scaled, using the SCTransform function in Seurat. Linear dimensionality reduction was then performed on both GEX and FB data, followed by weighted nearest neighbors clustering on the basis of the first 35 PCs for GEX and the first 10 PCs for FB using a resolution of 0.5, allowing identification of distinct cell states while incorporating both transcriptional and protein-level data ^77^. Cell cycle scores were calculated using the CellCycleScoring function in Seurat. Differentially expressed genes were calculated by Wilcoxon rank-sum test. Chi-squared analysis to determine sample enrichment within each cluster was performed via the chisq.test R function.

### *In vitro* model generation and CAR-T assay

Microtumors were generated according to Lam et al ^58^. Briefly, 1.5×10^4^ U87 MG IL13Rα2^+^ RFP^+^ cells were seeded in a hanging drop and allowed to form dense spheroids over 4 days before resuspending in a fibrin gel with a mix of human primary brain endothelial cells, pericytes and astrocytes, and injected into the central well of the OrganiX microfluidic device (AIM biotech). Over 7 days, a perfusable vasculature formed around the tumor spheroid, and the tumor cells invade into the perivascular space. 1×10^4^ transduced CAR 4 or 13BBζ were introduced into one side channel of the device, and microtumors with no CAR-T were used as control. 5 IU/ml IL-2 was supplemented in the media, and 50 μl of media was topped up every 3 days. Tumor spheroids and CAR-T infiltration were imaged on a Zeiss LSM880 inverted microscope with an environmental chamber and a 10X lens just before CAR-T addition (day 0) and on days 3, 6 and 9. Images were imported into <u>FIJI</u> and radial signal intensities were measured using the <u>ImageJ</u> built-in <u>Radial Plot plugin</u>.

### Xenogeneic mouse models

All animal studies were performed in accordance with guidelines approved by the MIT Division of Comparative Medicine and MIT Committee on Animal Care (Institutional Animal Care and Use Committee, protocol number 0621-032-24). Male NOD/SCID/IL2Rnull (NSG) mice were purchased from Jackson Laboratory and housed in the animal facilities at MIT. For *in vivo* studies, 6 to 8-week-told mice weighing between 24 and 31 g were injected subcutaneously on the right flank with 2×10^6^ IL13Rα2^+^ U87 cells, followed 21 days later by intravenous administration of 1×10^6^ CD3^+^ CAR-T cells or untransduced T cells, which were prepared as described above, via the tail vein. Tumor progression was subsequently monitored every 2-3 days using caliper measurement and the IVIS Spectrum imaging system (PerkinElmer) to measure bioluminescent signal after intraperitoneal administration of 0.15 mg of luciferin substrate per gram of body weight (PerkinElmer 122799). Total photon counts were quantified using LivingImage software. Mice were weighed every 2-3 days for signs of toxicity. Mice were euthanized upon observing signs of discomfort, morbidity, or limited mobility, or upon tumor area reaching 100 cm^2^ for untreated mice and 300 cm^2^ for CAR-treated mice.

### Statistical analysis

Statistical analyses were performed using the Prism (ver. 10) software, with the exception of the single-cell sequencing data, which were analyzed in R Studio using base packages or those described above. For microphysiological *in vitro* models, signal intensities were exported to Prism (ver. 10.1.1) and area under the curves were calculated. Sample sizes were not pre-determined using statistical methods. For statistical comparisons between multiple groups vs. control, significance was determined using two-way analysis of variance (ANOVA). Adjusted P values < 0.05 after multiple hypothesis correction with Dunnett’s multiple comparisons correction were considered statistically significant. The statistical test used for each experiment is noted in the relevant figure legend.

## DATA AVAILABILITY

### Data and materials availability

The NGS selection datasets have been deposited in the Sequence Read Archive and are available under the accession number PRJNA957857. The scRNA-seq data have been deposited in the Gene Expression Omnibus under accession number GSE259352. All data generated or analyzed during the study (including the DomainSeq-processed CARPOOL selection data) are included in the paper or its supplementary information.

### Code availability

The code used to analyze the domain composition of selected CARs can be accessed in the DomainSeq repository at https://github.com/birnbaumlab/Gordon-et-al-2022.

## ACKNOWLEDGEMENTS

We thank the Koch Institute’s Robert A. Swanson (1969) Biotechnology Center for their technical support, especially the Flow Cytometry Facility, Preclinical Modeling, Imaging and Testing Core, MIT BioMicro Center and High Throughput Sciences Core. We thank G. Paradis, P. Chamberlain, H. Holcombe, V. Spanoudaki and S. Levine for many helpful discussions and suggestions. M.E.B. was supported by a Packard Fellowship, a Pew-Stewart Scholarship and a grant from the Deshpande Center. D.A.L. was supported by US Army Research Office Cooperative Agreement W911NF-19-2-0026 Institute for Collaborative Biotechnologies. This work was supported in part by the Koch Institute Frontier Research Program (to M.E.B.), and the Koch Institute Support (core) Grant P30-CA14051 from the National Cancer Institute. This work was additionally supported by National Science Foundation Graduate Research Fellowships awarded to K.S.G., C.R.P., and A.D., by the Ludwig Graduate Fellowship awarded to A.D., and by award number T32GM144273 from the National Institute of General Medical Sciences awarded to A.G. The content is solely the responsibility of the authors and does not necessarily represent the official views of the National Institute of General Medical Sciences or the National Institutes of Health. This research is supported in part by the National Research Foundation, Prime Minister’s Office, Singapore under its Campus for Research Excellence and Technological Enterprise (CREATE) programme, through Singapore MIT Alliance for Research and Technology (SMART): Critical Analytics for Manufacturing Personalised-Medicine (CAMP) Inter-Disciplinary Research Group. We thank the Institute of A*STAR Cell and Molecular Biology (IMCB) Central Imaging Facility and flow cytometry unit. This research was also supported by the A*STAR Career Development Fund (C210112058) to M.S.Y.L and A*STAR core funding to A.P.

## AUTHOR CONTRIBUTIONS

Conceptualization: KSG, MEB

Methodology: KSG, CRP, MSYL

Investigation: KSG, CRP, AG, AD, MSYL, JJA

Visualization: KSG, CRP, AG, MSYL

Funding acquisition: MEB, AP, DAL

Supervision: MEB, AP, DAL

Writing – original draft: KSG

Writing – review & editing: KSG, CRP, MSYL, AG, AD, DAL, AP, MEB

## DECLARATION OF INTERESTS

The library approach described in this paper is the subject of a US patent application (PCT/US2020/017794) with M.E.B. as an inventor. A.P. is a member of the scientific advisory board and equity holder of AIM Biotech Pte. Ltd. M.E.B. is a founder, consultant, and equity holder of Kelonia Therapeutics and Abata Therapeutics. Khloe Gordon is currently employed and holds equity at Ginkgo Bioworks, Inc. The other authors declare no competing interests.

## Notes

### Summary of Updates

Corrected errors in method section - ensured figures show individual donors for comparison between CARs unless otherwise noted.

## REFERENCES

1. Lim, M., Xia, Y., Bettegowda, C., and Weller, M. (2018). Current state of immunotherapy for glioblastoma. Nat. Rev. Clin. Oncol. 15, 422–442.

2. Abbott, N.J., Rönnbäck, L., and Hansson, E. (2006). Astrocyte-endothelial interactions at the blood-brain barrier. Nat. Rev. Neurosci. 7, 41–53.

3. Terstappen, G.C., Meyer, A.H., Bell, R.D., and Zhang, W. (2021). Strategies for delivering therapeutics across the blood-brain barrier. Nat. Rev. Drug Discov. 20, 362–383.

4. Nance, E., Pun, S.H., Saigal, R., and Sellers, D.L. (2021). Drug delivery to the central nervous system. Nat. Rev. Mater. 7, 314–331.

5. Galea, I., Bernardes-Silva, M., Forse, P.A., van Rooijen, N., Liblau, R.S., and Perry, V.H. (2007). An antigen-specific pathway for CD8 T cells across the blood-brain barrier. J. Exp. Med. 204, 2023–2030.

6. Calzascia, T., Di Berardino-Besson, W., Wilmotte, R., Masson, F., de Tribolet, N., Dietrich, P.-Y., and Walker, P.R. (2003). Cutting edge: cross-presentation as a mechanism for efficient recruitment of tumor-specific CTL to the brain. J. Immunol. 171, 2187–2191.

7. Fraietta, J.A., Lacey, S.F., Orlando, E.J., Pruteanu-Malinici, I., Gohil, M., Lundh, S., Boesteanu, A.C., Wang, Y., O’Connor, R.S., Hwang, W.-T., et al. (2018). Determinants of response and resistance to CD19 chimeric antigen receptor (CAR) T cell therapy of chronic lymphocytic leukemia. Nat. Med. 24, 563–571.

8. Sabatino, M., Hu, J., Sommariva, M., Gautam, S., Fellowes, V., Hocker, J.D., Dougherty, S., Qin, H., Klebanoff, C.A., Fry, T.J., et al. (2016). Generation of clinical-grade CD19-specific CAR-modified CD8+ memory stem cells for the treatment of human B-cell malignancies. Blood 128, 519–528.

9. Gattinoni, L., Lugli, E., Ji, Y., Pos, Z., Paulos, C.M., Quigley, M.F., Almeida, J.R., Gostick, E., Yu, Z., Carpenito, C., et al. (2011). A human memory T cell subset with stem cell–like properties. Nat. Med. 17, 1290–1297.

10. Melenhorst, J.J., Chen, G.M., Wang, M., Porter, D.L., Chen, C., Collins, M.A., Gao, P., Bandyopadhyay, S., Sun, H., Zhao, Z., et al. (2022). Decade-long leukaemia remissions with persistence of CD4+ CAR T cells. Nature 602, 503–509.

11. Brown, C.E., Alizadeh, D., Starr, R., Weng, L., Wagner, J.R., Naranjo, A., Ostberg, J.R., Blanchard, M.S., Kilpatrick, J., Simpson, J., et al. (2016). Regression of Glioblastoma after Chimeric Antigen Receptor T-Cell Therapy. N. Engl. J. Med. 375, 2561–2569.

12. Sampson, J.H., Choi, B.D., Sanchez-Perez, L., Suryadevara, C.M., Snyder, D.J., Flores, C.T., Schmittling, R.J., Nair, S.K., Reap, E.A., Norberg, P.K., et al. (2014). EGFRvIII mCAR-modified T-cell therapy cures mice with established intracerebral glioma and generates host immunity against tumor-antigen loss. Clin. Cancer Res. 20, 972–984.

13. Adachi, K., Kano, Y., Nagai, T., Okuyama, N., Sakoda, Y., and Tamada, K. (2018). IL-7 and CCL19 expression in CAR-T cells improves immune cell infiltration and CAR-T cell survival in the tumor. Nat. Biotechnol. 36, 346–351.

14. Lai, J., Mardiana, S., House, I.G., Sek, K., Henderson, M.A., Giuffrida, L., Chen, A.X.Y., Todd, K.L., Petley, E.V., Chan, J.D., et al. (2020). Adoptive cellular therapy with T cells expressing the dendritic cell growth factor Flt3L drives epitope spreading and antitumor immunity. Nat. Immunol. 21, 914–926.

15. Choi, B.D., Yu, X., Castano, A.P., Bouffard, A.A., Schmidts, A., Larson, R.C., Bailey, S.R., Boroughs, A.C., Frigault, M.J., Leick, M.B., et al. (2019). CAR-T cells secreting BiTEs circumvent antigen escape without detectable toxicity. Nat. Biotechnol. 37, 1049–1058.

16. Yin, Y., Rodriguez, J.L., Li, N., Thokala, R., Nasrallah, M.P., Hu, L., Zhang, L., Zhang, J.V., Logun, M.T., Kainth, D., et al. (2022). Locally secreted BiTEs complement CAR T cells by enhancing killing of antigen heterogeneous solid tumors. Mol. Ther. 30, 2537–2553.

17. Majzner, R.G., and Mackall, C.L. (2019). Clinical lessons learned from the first leg of the CAR T cell journey. Nat. Med. 25, 1341–1355.

18. Maude, S.L., Laetsch, T.W., Buechner, J., Rives, S., Boyer, M., Bittencourt, H., Bader, P., Verneris, M.R., Stefanski, H.E., Myers, G.D., et al. (2018). Tisagenlecleucel in children and young adults with B-cell lymphoblastic leukemia. N. Engl. J. Med. 378, 439–448.

19. Locke, F.L., Miklos, D.B., Jacobson, C.A., Perales, M.-A., Kersten, M.-J., Oluwole, O.O., Ghobadi, A., Rapoport, A.P., McGuirk, J., Pagel, J.M., et al. (2022). Axicabtagene ciloleucel as second-line therapy for large B-cell lymphoma. N. Engl. J. Med. 386, 640–654.

20. Schuster, S.J., Bishop, M.R., Tam, C.S., Waller, E.K., Borchmann, P., McGuirk, J.P., Jäger, U., Jaglowski, S., Andreadis, C., Westin, J.R., et al. (2019). Tisagenlecleucel in adult relapsed or refractory diffuse large B-cell lymphoma. N. Engl. J. Med. 380, 45–56.

21. Morgan, R.A., Yang, J.C., Kitano, M., Dudley, M.E., Laurencot, C.M., and Rosenberg, S.A. (2010). Case report of a serious adverse event following the administration of T cells transduced with a chimeric antigen receptor recognizing ERBB2. Mol. Ther. 18, 843–851.

22. Maggs, L., Cattaneo, G., Dal, A.E., Moghaddam, A.S., and Ferrone, S. (2021). CAR T cell-based immunotherapy for the treatment of glioblastoma. Front. Neurosci. 15, 662064.

23. Johnson, L.A., Scholler, J., Ohkuri, T., Kosaka, A., Patel, P.R., McGettigan, S.E., Nace, A.K., Dentchev, T., Thekkat, P., Loew, A., et al. (2015). Rational development and characterization of humanized anti-EGFR variant III chimeric antigen receptor T cells for glioblastoma. Sci. Transl. Med. 7, 275ra22.

24. Durgin, J.S., Henderson, F., Jr, Nasrallah, M.P., Mohan, S., Wang, S., Lacey, S.F., Melenhorst, J.J., Desai, A.S., Lee, J.Y.K., Maus, M.V., et al. (2021). Case report: Prolonged survival following EGFRvIII CAR T cell treatment for recurrent glioblastoma. Front. Oncol. 11, 669071.

25. Wang, D., Starr, R., Chang, W.-C., Aguilar, B., Alizadeh, D., Wright, S.L., Yang, X., Brito, A., Sarkissian, A., Ostberg, J.R., et al. (2020). Chlorotoxin-directed CAR T cells for specific and effective targeting of glioblastoma. Sci. Transl. Med. 12, eaaw2672.

26. Prapa, M., Chiavelli, C., Golinelli, G., Grisendi, G., Bestagno, M., Di Tinco, R., Dall’Ora, M., Neri, G., Candini, O., Spano, C., et al. (2021). GD2 CAR T cells against human glioblastoma. NPJ Precis. Oncol. 5, 93.

27. Majzner, R.G., Ramakrishna, S., Yeom, K.W., Patel, S., Chinnasamy, H., Schultz, L.M., Richards, R.M., Jiang, L., Barsan, V., Mancusi, R., et al. (2022). GD2-CAR T cell therapy for H3K27M-mutated diffuse midline gliomas. Nature 603, 934–941.

28. Tang, X., Zhao, S., Zhang, Y., Wang, Y., Zhang, Z., Yang, M., Zhu, Y., Zhang, G., Guo, G., Tong, A., et al. (2019). B7-H3 as a novel CAR-T therapeutic target for glioblastoma. Mol. Ther. Oncolytics 14, 279–287.

29. Tang, X., Wang, Y., Huang, J., Zhang, Z., Liu, F., Xu, J., Guo, G., Wang, W., Tong, A., and Zhou, L. (2021). Administration of B7-H3 targeted chimeric antigen receptor-T cells induce regression of glioblastoma. Signal Transduct. Target. Ther. 6, 125.

30. Harrer, D.C., Dörrie, J., and Schaft, N. (2019). CSPG4 as target for CAR-T-cell therapy of various tumor entities-merits and challenges. Int. J. Mol. Sci. 20, 5942.

31. Choe, J.H., Watchmaker, P.B., Simic, M.S., Gilbert, R.D., Li, A.W., Krasnow, N.A., Downey, K.M., Yu, W., Carrera, D.A., Celli, A., et al. (2021). SynNotch-CAR T cells overcome challenges of specificity, heterogeneity, and persistence in treating glioblastoma. Sci. Transl. Med. 13. 10.1126/scitranslmed.abe7378.

32. MacKay, M., Afshinnekoo, E., Rub, J., Hassan, C., Khunte, M., Baskaran, N., Owens, B., Liu, L., Roboz, G.J., Guzman, M.L., et al. (2020). The therapeutic landscape for cells engineered with chimeric antigen receptors. Nat. Biotechnol. 38, 233–244.

33. Gordon, K.S., Kyung, T., Perez, C.R., Holec, P.V., Ramos, A., Zhang, A.Q., Agarwal, Y., Liu, Y., Koch, C.E., Starchenko, A., et al. (2022). Screening for CD19-specific chimaeric antigen receptors with enhanced signalling via a barcoded library of intracellular domains. Nat. Biomed. Eng. 6, 855–866.

34. Goodman, D.B., Azimi, C.S., Kearns, K., Talbot, A., Garakani, K., Garcia, J., Patel, N., Hwang, B., Lee, D., Park, E., et al. (2022). Pooled screening of CAR T cells identifies diverse immune signaling domains for next-generation immunotherapies. Sci. Transl. Med. 14, eabm1463.

35. Daniels, K.G., Wang, S., Simic, M.S., Bhargava, H.K., Capponi, S., Tonai, Y., Yu, W., Bianco, S., and Lim, W.A. (2022). Decoding CAR T cell phenotype using combinatorial signaling motif libraries and machine learning. Science 378, 1194–1200.

36. Castellanos-Rueda, R., Di Roberto, R.B., Bieberich, F., Schlatter, F.S., Palianina, D., Nguyen, O.T.P., Kapetanovic, E., Läubli, H., Hierlemann, A., Khanna, N., et al. (2022). speedingCARs: accelerating the engineering of CAR T cells by signaling domain shuffling and single-cell sequencing. Nat. Commun. 13, 6555.

37. Castellanos-Rueda, R., Wang, K.-L.K., Forster, J.L., Driessen, A., Frank, J.A., Martínez, M.R., and Reddy, S.T. (2024). Dissecting the role of CAR signaling architectures on T cell activation and persistence using pooled screening and single-cell sequencing. bioRxiv. 10.1101/2024.02.26.582129.

38. Rios, X., Pardias, O., Morales, M.A., Bhattacharya, P., Chen, Y., Guo, L., Zhang, C., Di Pierro, E.J., Tian, G., Barragan, G.A., et al. (2023). Refining chimeric antigen receptors via barcoded protein domain combination pooled screening. Mol. Ther. 31, 3210–3224.

39. Brown, C.E., Hibbard, J.C., Alizadeh, D., Blanchard, M.S., Natri, H.M., Wang, D., Ostberg, J.R., Aguilar, B., Wagner, J.R., Paul, J.A., et al. (2024). Locoregional delivery of IL-13Rα2-targeting CAR-T cells in recurrent high-grade glioma: a phase 1 trial. Nat. Med., 1–12.

40. Okamoto, H., Yoshimatsu, Y., Tomizawa, T., Kunita, A., Takayama, R., Morikawa, T., Komura, D., Takahashi, K., Oshima, T., Sato, M., et al. (2019). Interleukin-13 receptor α2 is a novel marker and potential therapeutic target for human melanoma. Sci. Rep. 9, 1–13.

41. Kawakami, K., Kawakami, M., and Puri, R.K. (2004). Specifically targeted killing of interleukin-13 (IL-13) receptor-expressing breast cancer by IL-13 fusion cytotoxin in animal model of human disease. Mol. Cancer Ther. 3, 137–147.

42. Fujisawa, T., Joshi, B., Nakajima, A., and Puri, R.K. (2009). A novel role of interleukin-13 receptor alpha2 in pancreatic cancer invasion and metastasis. Cancer Res. 69, 8678–8685.

43. Nguyen, V., Conyers, J.M., Zhu, D., Gibo, D.M., Dorsey, J.F., Debinski, W., and Mintz, A. (2011). IL-13Rα2-Targeted Therapy Escapees: Biologic and Therapeutic Implications. Transl. Oncol. 4, 390–400.

44. Saikali, S., Avril, T., Collet, B., Hamlat, A., Bansard, J.-Y., Drenou, B., Guegan, Y., and Quillien, V. (2006). Expression of nine tumour antigens in a series of human glioblastoma multiforme: interest of EGFRvIII, IL-13Rα2, gp100 and TRP-2 for immunotherapy. Preprint, 10.1007/s11060-006-9220-3 10.1007/s11060-006-9220-3.

45. Newman, J.P., Wang, G.Y., Arima, K., Guan, S.P., Waters, M.R., Cavenee, W.K., Pan, E., Aliwarga, E., Chong, S.T., Kok, C.Y.L., et al. (2017). Interleukin-13 receptor alpha 2 cooperates with EGFRvIII signaling to promote glioblastoma multiforme. Nat. Commun. 8, 1913.

46. Jonnalagadda, M., Mardiros, A., Urak, R., Wang, X., Hoffman, L.J., Bernanke, A., Chang, W.-C., Bretzlaff, W., Starr, R., Priceman, S., et al. (2015). Chimeric antigen receptors with mutated IgG4 Fc spacer avoid fc receptor binding and improve T cell persistence and antitumor efficacy. Mol. Ther. 23, 757–768.

47. Lanier, L.L. (2006). Viral immunoreceptor tyrosine-based activation motif (ITAM)-mediated signaling in cell transformation and cancer. Trends Cell Biol. 16, 388–390.

48. Geginat, J., Sallusto, F., and Lanzavecchia, A. (2001). Cytokine-driven proliferation and differentiation of human naive, central memory, and effector memory CD4(+) T cells. J. Exp. Med. 194, 1711–1719.

49. Kagoya, Y., Tanaka, S., Guo, T., Anczurowski, M., Wang, C.-H., Saso, K., Butler, M.O., Minden, M.D., and Hirano, N. (2018). A novel chimeric antigen receptor containing a JAK-STAT signaling domain mediates superior antitumor effects. Nat. Med. 24, 352–359.

50. Wang, D., Aguilar, B., Starr, R., Alizadeh, D., Brito, A., Sarkissian, A., Ostberg, J.R., Forman, S.J., and Brown, C.E. (2018). Glioblastoma-targeted CD4+ CAR T cells mediate superior antitumor activity. JCI Insight 3. 10.1172/jci.insight.99048.

51. Han, J., Alvarez-Breckenridge, C.A., Wang, Q.-E., and Yu, J. (2015). TGF-β signaling and its targeting for glioma treatment. Am. J. Cancer Res. 5, 945–955.

52. Tran, T.-T., Uhl, M., Ma, J.Y., Janssen, L., Sriram, V., Aulwurm, S., Kerr, I., Lam, A., Webb, H.K., Kapoun, A.M., et al. (2007). Inhibiting TGF-beta signaling restores immune surveillance in the SMA-560 glioma model. Neuro. Oncol. 9, 259–270.

53. Salter, A.I., Ivey, R.G., Kennedy, J.J., Voillet, V., Rajan, A., Alderman, E.J., Voytovich, U.J., Lin, C., Sommermeyer, D., Liu, L., et al. (2018). Phosphoproteomic analysis of chimeric antigen receptor signaling reveals kinetic and quantitative differences that affect cell function. Sci. Signal. 11. 10.1126/scisignal.aat6753.

54. Frey, N., and Porter, D. (2019). Cytokine release syndrome with chimeric antigen receptor T cell therapy. Biol. Blood Marrow Transplant. 25, e123–e127.

55. Wei, J., Liu, Y., Wang, C., Zhang, Y., Tong, C., Dai, G., Wang, W., Rasko, J.E.J., Melenhorst, J.J., Qian, W., et al. (2020). The model of cytokine release syndrome in CAR T-cell treatment for B-cell non-Hodgkin lymphoma. Signal Transduct. Target. Ther. 5, 134.

56. Lu, K.H.-N., Michel, J., Kilian, M., Aslan, K., Qi, H., Kehl, N., Jung, S., Sanghvi, K., Lindner, K., Zhang, X.-W., et al. (2022). T cell receptor dynamic and transcriptional determinants of T cell expansion in glioma-infiltrating T cells. Neurooncol. Adv. 4, vdac140.

57. Szabo, P.A., Levitin, H.M., Miron, M., Snyder, M.E., Senda, T., Yuan, J., Cheng, Y.L., Bush, E.C., Dogra, P., Thapa, P., et al. (2019). Single-cell transcriptomics of human T cells reveals tissue and activation signatures in health and disease. Nat. Commun. 10, 4706.

58. Lam, M.S.Y., Aw, J.J.Y., Tan, D., Vijayakumar, R., Lim, H.Y.G., Yada, S., Pang, Q.Y., Barker, N., Tang, C., Ang, B.T., et al. (2023). Unveiling the influence of tumor microenvironment and spatial heterogeneity on temozolomide resistance in glioblastoma using an advanced human in vitro model of the bloodLJbrain barrier and glioblastoma. Small 19. 10.1002/smll.202302280.

59. Adriani, G., and Pavesi, A. (2024). The OrganiX microfluidic system to recreate the complex tumour microenvironment. Nat. Rev. Immunol., 1–1.

60. Schmidts, A., Srivastava, A.A., Ramapriyan, R., Bailey, S.R., Bouffard, A.A., Cahill, D.P., Carter, B.S., Curry, W.T., Dunn, G.P., Frigault, M.J., et al. (2023). Tandem chimeric antigen receptor (CAR) T cells targeting EGFRvIII and IL-13Rα2 are effective against heterogeneous glioblastoma. Neurooncol. Adv. 5. 10.1093/noajnl/vdac185.

61. Bagley, S.J., Logun, M., Fraietta, J.A., Wang, X., Desai, A.S., Bagley, L.J., Nabavizadeh, A., Jarocha, D., Martins, R., Maloney, E., et al. (2024). Intrathecal bivalent CAR T cells targeting EGFR and IL13Rα2 in recurrent glioblastoma: phase 1 trial interim results. Nat. Med., 1–10.

62. Choi, B.D., Gerstner, E.R., Frigault, M.J., Leick, M.B., Mount, C.W., Balaj, L., Nikiforow, S., Carter, B.S., Curry, W.T., Gallagher, K., et al. (2024). Intraventricular CARv3-TEAM-E T cells in recurrent glioblastoma. N. Engl. J. Med. 10.1056/nejmoa2314390.

63. Feucht, J., Sun, J., Eyquem, J., Ho, Y.-J., Zhao, Z., Leibold, J., Dobrin, A., Cabriolu, A., Hamieh, M., and Sadelain, M. (2019). Calibration of CAR activation potential directs alternative T cell fates and therapeutic potency. Preprint, 10.1038/s41591-018-0290-5 10.1038/s41591-018-0290-5.

64. Turnbull, I.R., and Colonna, M. (2007). Activating and inhibitory functions of DAP12. Nat. Rev. Immunol. 7, 155–161.

65. Sun, M., Xu, P., Wang, E., Zhou, M., Xu, T., Wang, J., Wang, Q., Wang, B., Lu, K., Wang, C., et al. (2021). Novel two-chain structure utilizing KIRS2/DAP12 domain improves the safety and efficacy of CAR-T cells in adults with r/r B-ALL. Mol. Ther. Oncolytics 23, 96– 106.

66. Zheng, L., Ren, L., Kouhi, A., Khawli, L.A., Hu, P., Kaslow, H.R., and Epstein, A.L. (2020). A humanized Lym-1 CAR with novel DAP10/DAP12 signaling domains demonstrates reduced tonic signaling and increased antitumor activity in B-cell lymphoma models. Clin. Cancer Res. 26, 3694–3706.

67. Jan, C.-I., Huang, S.-W., Canoll, P., Bruce, J.N., Lin, Y.-C., Pan, C.-M., Lu, H.-M., Chiu, S.-C., and Cho, D.-Y. (2021). Targeting human leukocyte antigen G with chimeric antigen receptors of natural killer cells convert immunosuppression to ablate solid tumors. J. Immunother. Cancer 9, e003050.

68. Ng, Y.-Y., Tay, J.C.K., Li, Z., Wang, J., Zhu, J., and Wang, S. (2021). T cells expressing NKG2D CAR with a DAP12 signaling domain stimulate lower cytokine production while effective in tumor eradication. Mol. Ther. 29, 75–85.

69. Töpfer, K., Cartellieri, M., Michen, S., Wiedemuth, R., Müller, N., Lindemann, D., Bachmann, M., Füssel, M., Schackert, G., and Temme, A. (2015). DAP12-based activating chimeric antigen receptor for NK cell tumor immunotherapy. J. Immunol. 194, 3201–3212.

70. Müller, N., Michen, S., Tietze, S., Töpfer, K., Schulte, A., Lamszus, K., Schmitz, M., Schackert, G., Pastan, I., and Temme, A. (2015). Engineering NK cells modified with an EGFRvIII-specific chimeric antigen receptor to overexpress CXCR4 improves immunotherapy of CXCL12/SDF-1α-secreting glioblastoma. J. Immunother. 38, 197–210.

71. Chen, B., Zhou, M., Zhang, H., Wang, C., Hu, X., Wang, B., and Wang, E. (2019). TREM1/Dap12-based CAR-T cells show potent antitumor activity. Immunotherapy 11, 1043–1055.

72. Chen, X., Bai, F., Sokol, L., Zhou, J., Ren, A., Painter, J.S., Liu, J., Sallman, D.A., Chen, Y.A., Yoder, J.A., et al. (2009). A critical role for DAP10 and DAP12 in CD8+ T cell-mediated tissue damage in large granular lymphocyte leukemia. Blood 113, 3226–3234.

73. Sedý, J.R., Spear, P.G., and Ware, C.F. (2008). Cross-regulation between herpesviruses and the TNF superfamily members. Nat. Rev. Immunol. 8, 861–873.

74. Bai, Z., Woodhouse, S., Zhao, Z., Arya, R., Govek, K., Kim, D., Lundh, S., Baysoy, A., Sun, H., Deng, Y., et al. (2022). Single-cell antigen-specific landscape of CAR T infusion product identifies determinants of CD19-positive relapse in patients with ALL. Sci. Adv. 8. 10.1126/sciadv.abj2820.

75. Stuart, T., Butler, A., Hoffman, P., Hafemeister, C., Papalexi, E., Mauck, W.M., 3rd, Hao, Y., Stoeckius, M., Smibert, P., and Satija, R. (2019). Comprehensive integration of single-cell data. Cell 177, 1888–1902.e21.

76. Stoeckius, M., Zheng, S., Houck-Loomis, B., Hao, S., Yeung, B.Z., Mauck, W.M., 3rd, Smibert, P., and Satija, R. (2018). Cell Hashing with barcoded antibodies enables multiplexing and doublet detection for single cell genomics. Genome Biol. 19, 224.

77. Hao, Y., Hao, S., Andersen-Nissen, E., Mauck, W.M., III, Zheng, S., Butler, A., Lee, M.J., Wilk, A.J., Darby, C., Zager, M., et al. (2021). Integrated analysis of multimodal single-cell data. Cell 184, 3573–3587.e29.

