## Supplemental Information for "Pooled screening for CAR function identifies novel IL13Rα2-targeted CARs for treatment of glioblastoma"

**List of Supplemental Materials**

Supplemental Figures 1-12

Supplemental Table 1

Supplemental Data File S1 to S2 (Excel files)

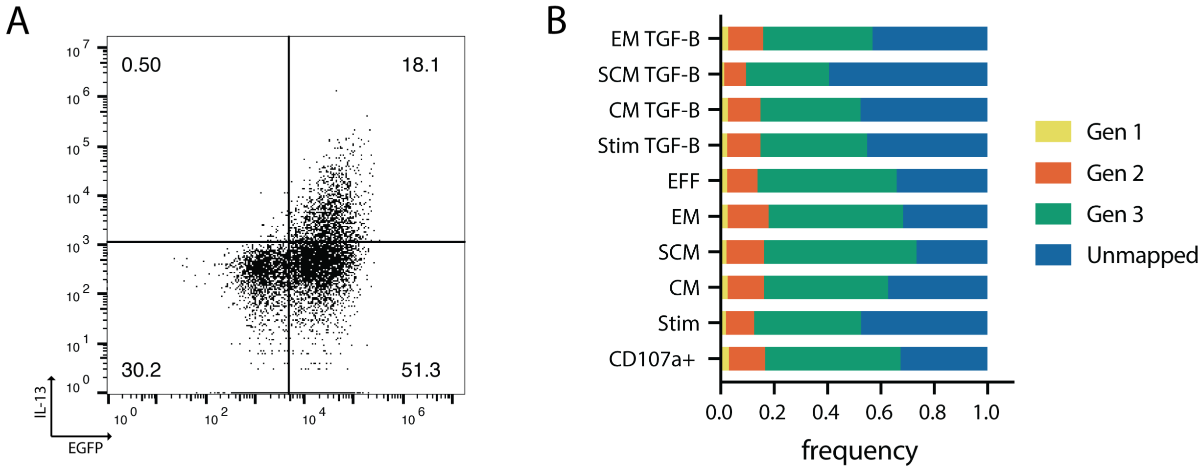


**Supplemental Figure 1**. Related to Figures 1-2. (**A**) FACS plots shown for transduced $CD4^+$ library populations prior to sorting for EGFP, with CAR expression measured by IL-13 expression (donor 1). (**B**) Distribution of CAR generations mapped to each selected population.

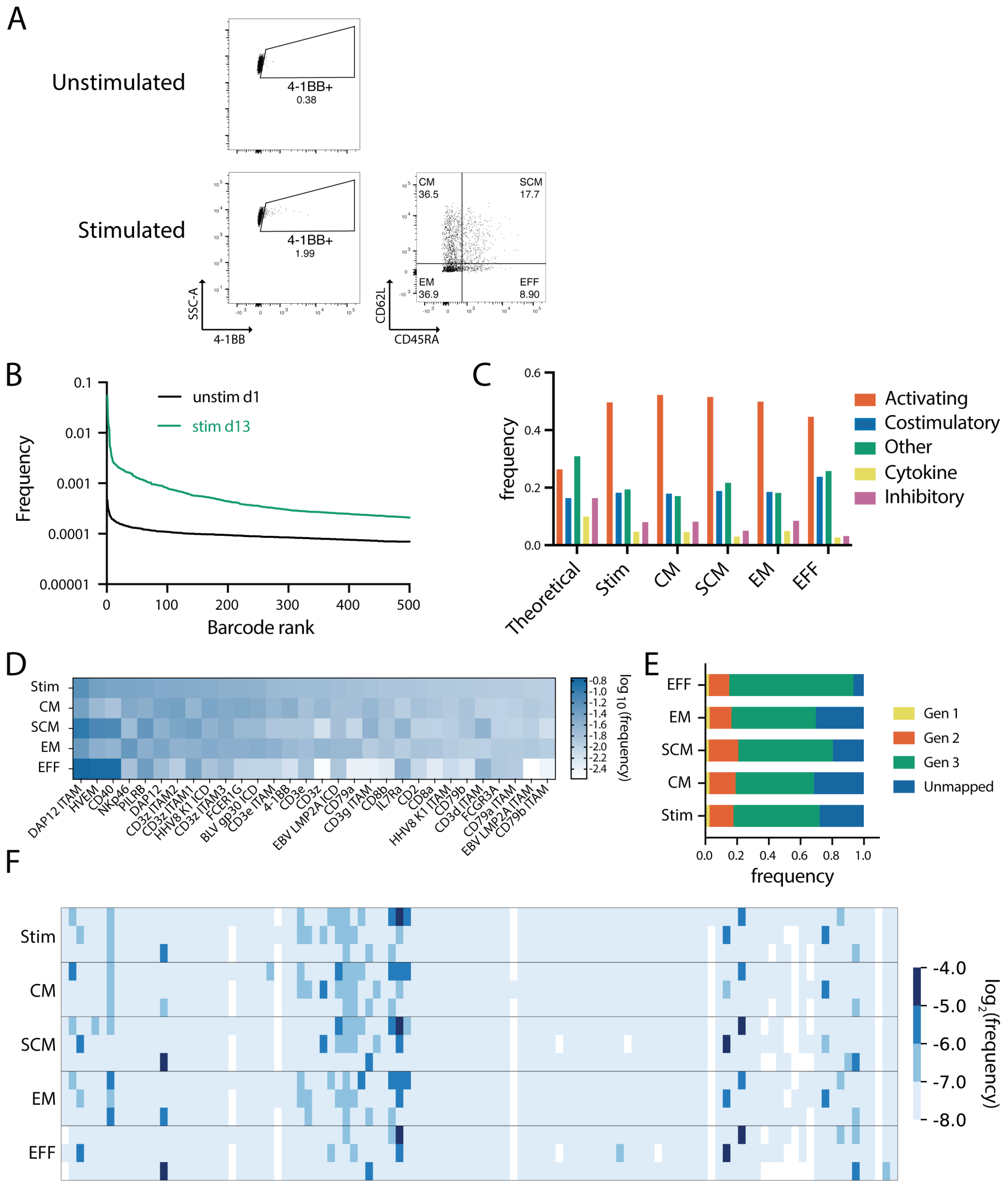


**Supplemental Figure 2.** $CD4^+$ biological replicate selection (donor 2). (**A**) FACS plots depicting sorted activation and memory populations. (**B**) Barcode enrichment for unselected and selected populations, with barcode frequency on the y-axis and barcode rank (by frequency) on the x-axis. (**C**) Frequency of each family of signaling domain throughout different selected populations. (**D**) Heat maps showing bulk ICD $log_{10}$ frequencies (irrespective of position) for rechallenged CARs. (**E**) Distribution of CAR generations mapped to each selected population. (**F**) $Log_2$ frequencies of ICDs at each intracellular position relative to the transmembrane domain for selected populations.

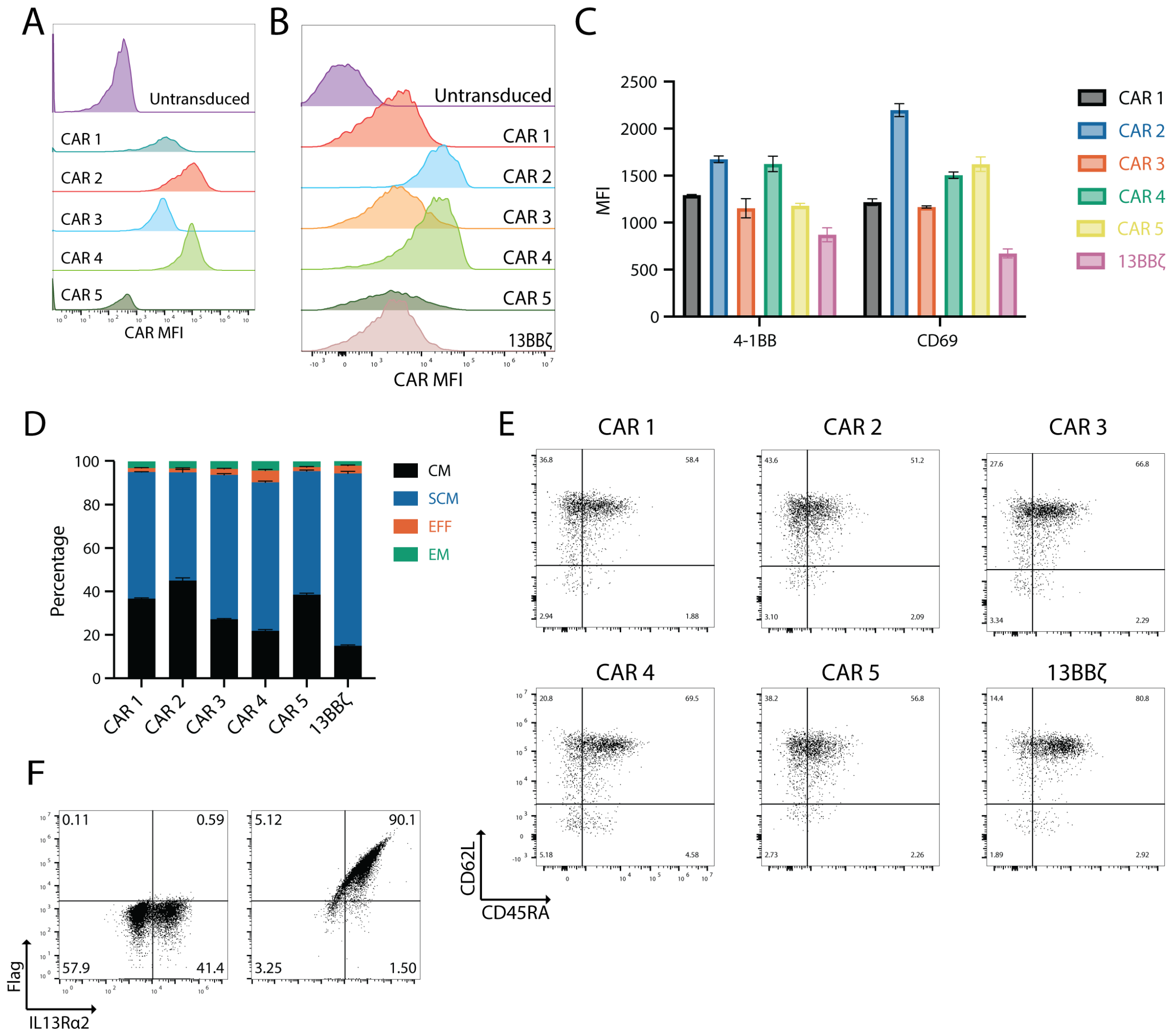


**Supplemental Figure 3.** Basal phenotypes of selected CARs. CAR expression, as measured by IL-13 staining, in (**A**) Jurkat and (**B**) human primary CD4$^{+}$ T cells for candidate CARs. (**C**) Basal CD69 and 4-1BB expression in unstimulated CD4$^{+}$ CARs. (**D**) Memory phenotype of unstimulated candidate CARs. (**E**) FACS plots showing memory phenotypes of unstimulated candidate CARs. (**F**) Surface expression of Flag and IL13Rα2 in wild-type (left) and engineered Flag-tagged IL13Rα2$^{+}$ (right) U87 cells. Panels (**C-E**) are representative of three biological replicates. *P* values in (**C**) relative to control CAR are 0.0001, <0.0001, 0.0095, <0.0001, and 0.0040 for CAR 1, 2, 3, 4, and 5 4-1BB expression, respectively. For CD69 expression*, P* values relative to control CAR are <0.0001 for all non-control CARs.

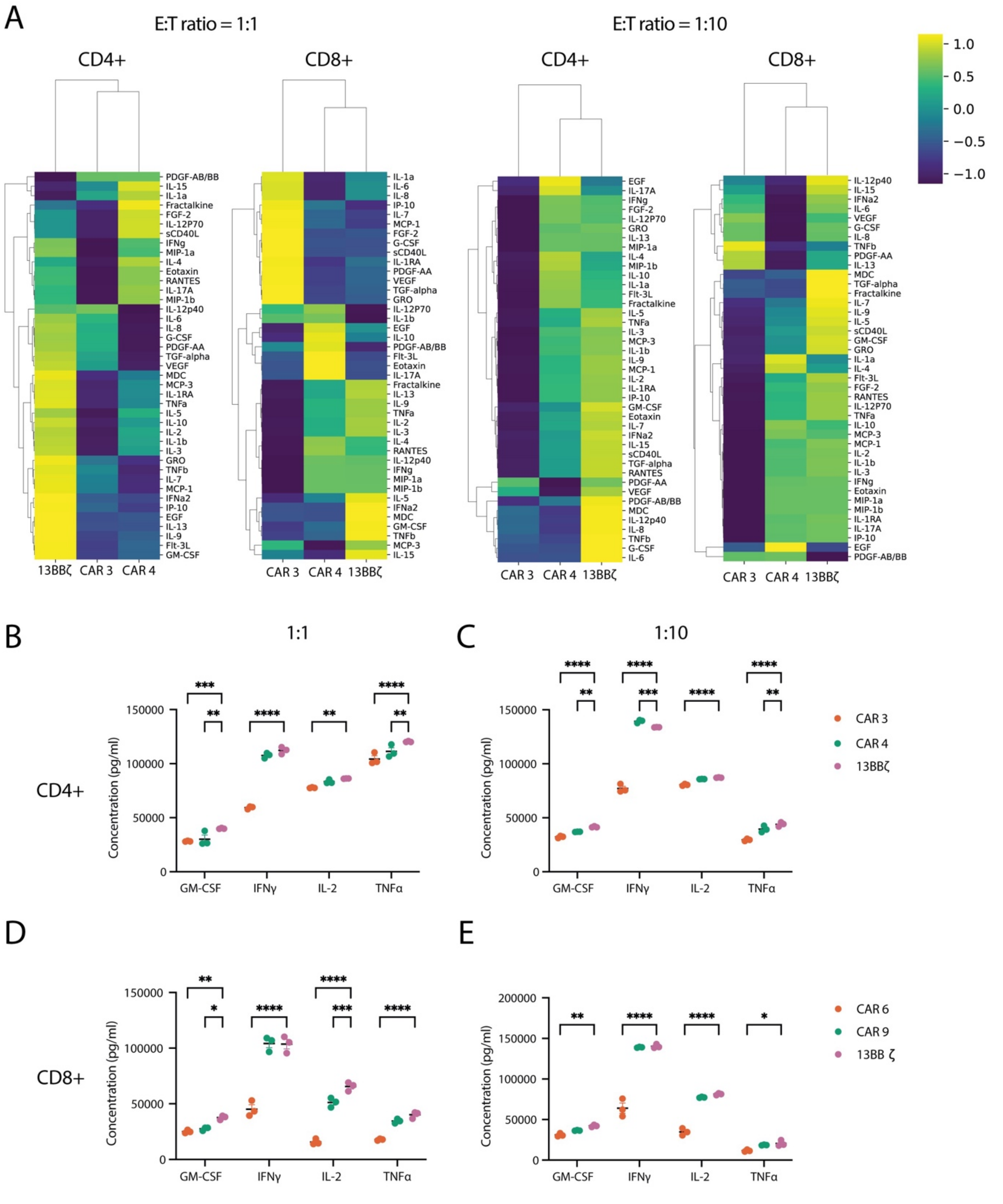


**Supplemental Figure 4.** (**A**) Cytokine secretion shown for human primary CD4$^+$ and CD8$^+$ T cells stimulated at a 1:1 or 1:10 E:T ratio with IL13Rα2$^+$ U87 cells. (**B-E**) Individual cytokine secretion levels plotted for GM-CSF, IFNγ, IL-2, IL-6, and TNFα for CD4$^+$ and CD8$^+$ T cells at E:T ratios of 1:1 and 1:10. Data in heat maps are z-scored. *P* values were calculated using two-way ANOVA with Dunnett's multiple comparisons test. In (**B**), they are 0.0004 for CAR 3 vs. 13BBζ and 0.0022 for CAR 4 vs. 13BBζ for GM-CSF. For IFNγ, the *P* value is <0.0001 for CAR 3 vs. 13BBζ. For

IL-2, the *P* value is 0.0077 for CAR 3 vs. 13BBζ. For TNFα, the *P* values are <0.0001 for CAR 3 vs. 13BBζ and 0.0045 for CAR 4 vs. 13BBζ. In (**C**), the *P* values are <0.0001 for CAR 3 vs. 13BBζ and 0.0041 for CAR 4 vs. 13BBζ for GM-CSF. For IFNγ, the *P* values are <0.0001 for CAR 3 vs. 13BBζ and 0.0008 for CAR 4 vs. 13BBζ. For IL-2, the *P* value is <0.0001 for CAR 3 vs. 13BBζ. For TNFα, the *P* values are <0.0001 for CAR 3 vs. 13BBζ and 0.0054 for CAR 4 vs. 13BBζ. In (**D**), the *P* values are 0.0021 for CAR 3 vs. 13BBζ and 0.0127 for CAR 4 vs. 13BBζ. For IFNγ and IL-2, the *P* values are <0.0001 for CAR 3 and 0.0006 for CAR 4 vs. 13BBζ IL-2 secretion. For TNFα, the *P* values is <0.0001 for CAR 3 vs. 13BBζ. In (**E**), the *P* values for GM-CSF are 0.0024 for CAR 3 vs. 13BBζ. For IFNγ, the *P* value is <0.0001 for CAR 3 vs. 13BBζ. For IL-2, the *P* values is <0.0001 for CAR 3 vs. 13BBζ . For TNFα, the *P* value is 0.0143 for CAR 3 vs. 13BBζ. For all data, n = 3 per group and df = 24. Representative of two biological replicates.

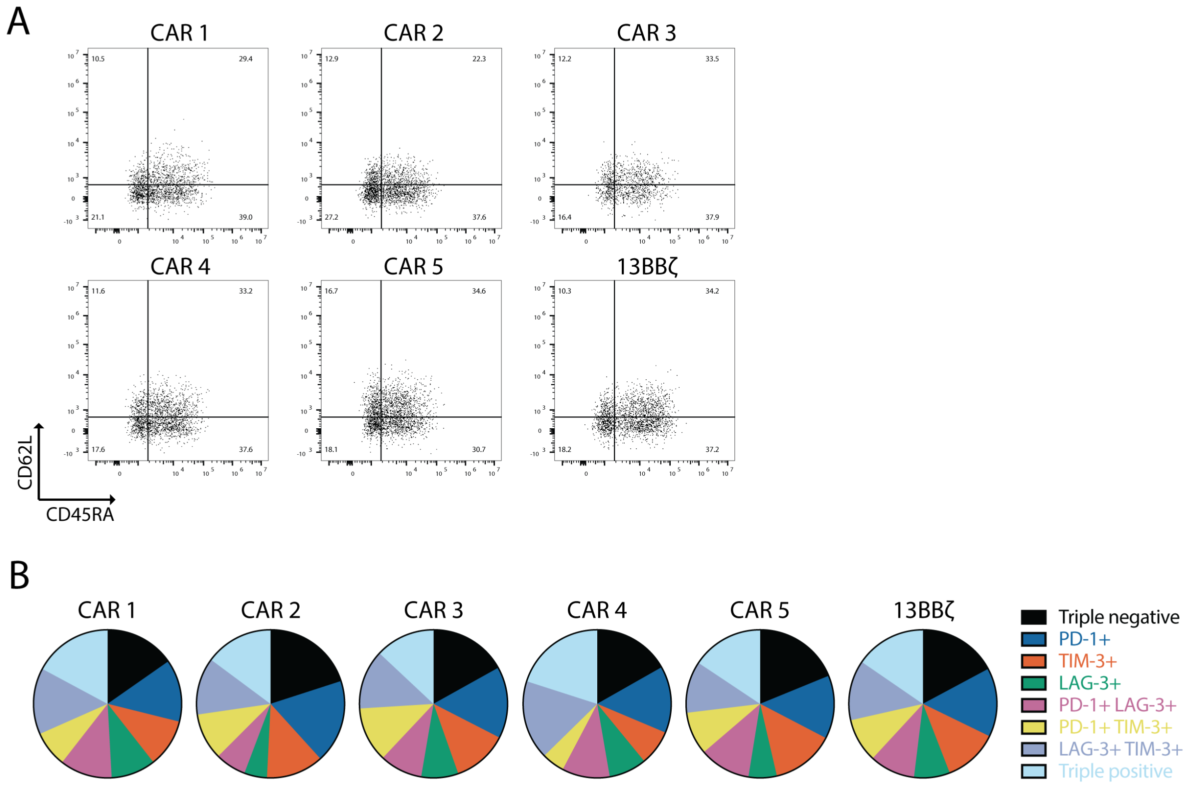


**Supplemental Figure 5.** Related to Figure 4. (**A**) FACS plots showing memory phenotypes on day 14 of rechallenge. (**B**) Co-expression of exhaustion markers on day 14 of rechallenge. Representative of two biological replicates.

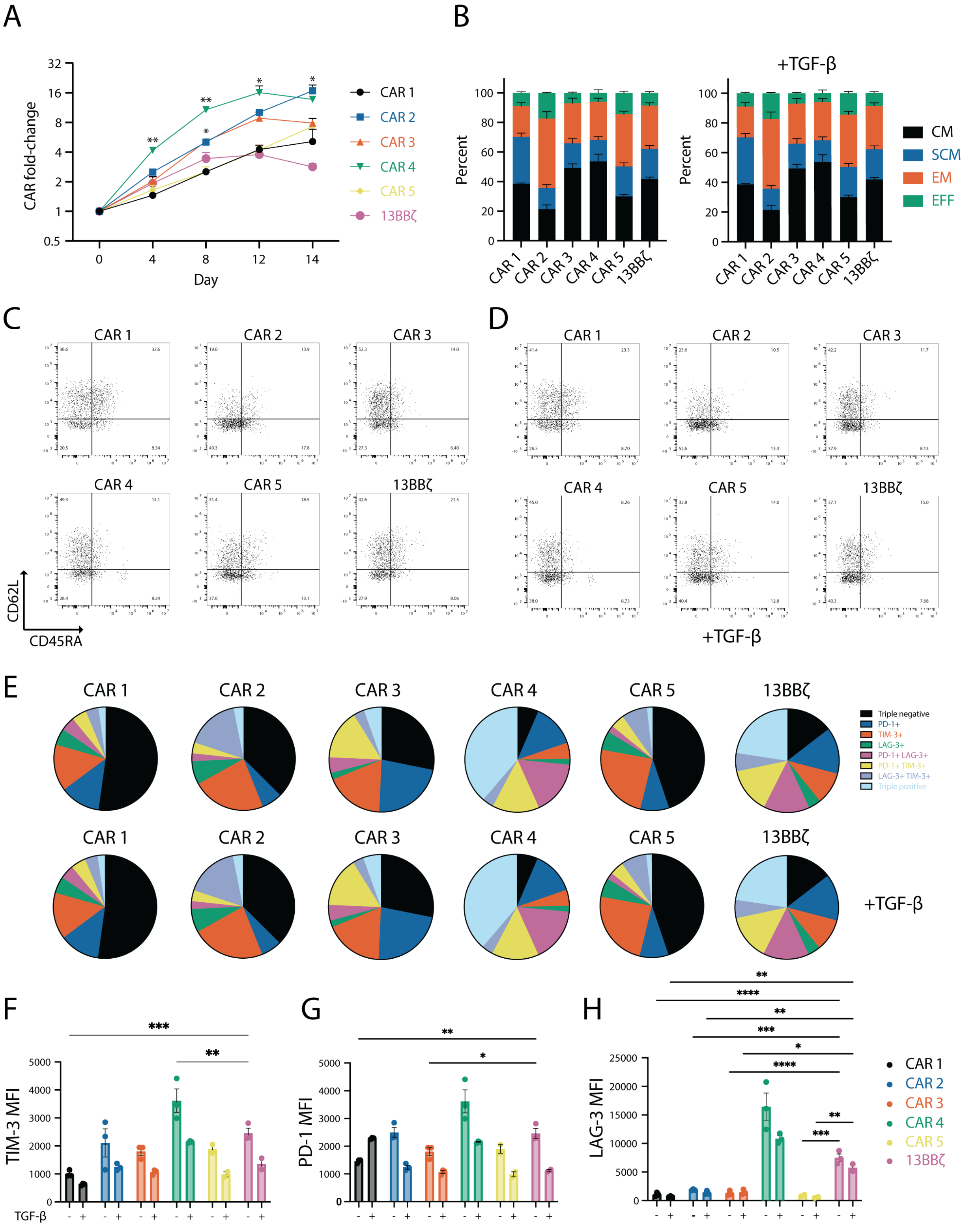


**Supplemental Figure 6.** Selected CARs persist and proliferate in response to tumor rechallenge with wild-type U87. (**A**) CAR fold-change following rechallenge with wild-type U87 cells at a 1:1 E:T ratio on days 0, 4, 8, and 12, with TGF-$\beta$ supplementation on days 8 and 12 at 5 ng/ml. (**B-D**)

Memory and (**E-H**) exhaustion phenotypes on day 14 of rechallenge. Data for (**A-B**) shows means ± s.e.m. (n = 3). $P$ values in (**A**) were determined using mixed-effects analysis with Dunnett's multiple comparisons test and are 0.0033 for CAR 4 vs. 13BB$\zeta$ on day 4, 0.0280 and 0.0073 for CAR 3 vs. 13BB$\zeta$ and CAR 4 vs. 13 BB$\zeta$ on day 8, 0.0182 for CAR 4 vs. 13BB$\zeta$ on day 12, and 0.0141 for CAR 4 vs. 13BB$\zeta$ on day 14 (n = 3, df = 2). Data is representative of two biological replicates. (**C,D**) FACS plots showing memory phenotype of rechallenged CARs on day 14 (**C**) without or (**D**) with TGF-$\beta$ treatment. Expression of exhaustion markers by human primary $CD4^+$ CAR-T cells on day 14 of rechallenge, as measured by (**F**) TIM-3, (**G**) PD-1, and (**H**) LAG-3 expression. $P$ values in (**F-H**) were determined using two-way ANOVA analysis with Dunnett's multiple comparisons test and, in (**H**), are 0.0007 for CAR 1 vs. 13BB$\zeta$ and 0.0059 for CAR 4 vs. 13BB$\zeta$ without TGF-$\beta$ treatment. In (**G**), $P$ values are 0.0012 for CAR 1 vs. 13BB$\zeta$ and 0.0398 for CAR 3 vs. 13BB$\zeta$ without TGF-$\beta$ treatment. In (**H**), $P$values are $<$0.0001 for CARs 1 and 3 vs. 13BB$\zeta$ and 0.0003 for CAR 2 vs. 13BB$\zeta$, and 0.0002 for CAR 5 vs. 13BB$\zeta$ without TGF-$\beta$ treatment; with TGF-$\beta$ treatment, $P$ values are 0.0026 for CAR 1 vs. 13BB$\zeta$, 0.0086 for CAR 2 vs. 13BB$\zeta$, 0.0106 for CAR 3 vs. 13BB$\zeta$, 0.0027 for CAR 4 vs. 13BB$\zeta$, and 0.0047 for CAR 5 vs. 13BB$\zeta$.

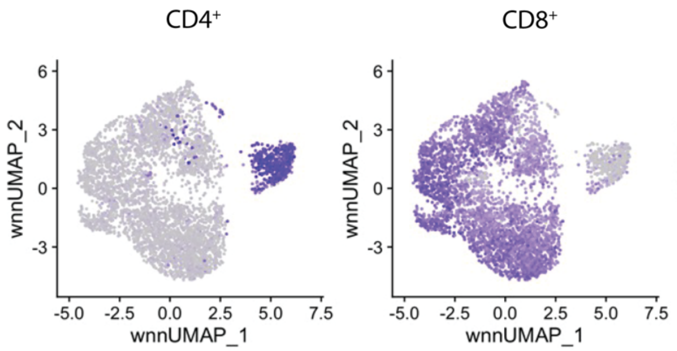


**Supplemental Figure 7.** Differential of CD4 (left) and CD8 (right) expression, as determined by single cell sequencing shown in Figure 5.

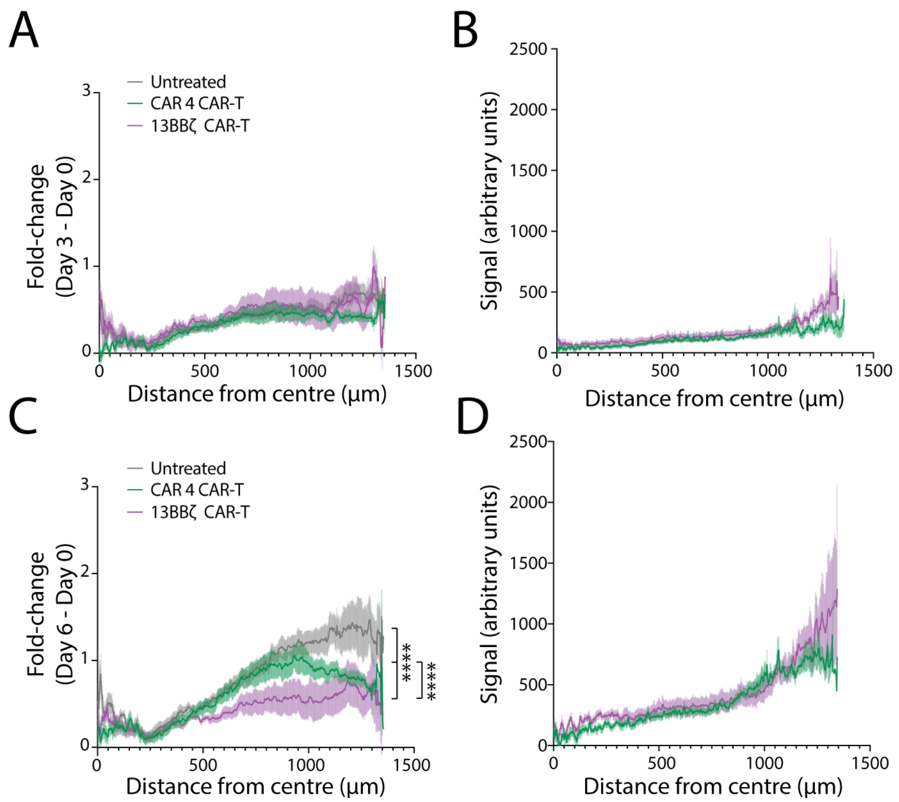


**Supplemental Figure 8.** Increase in U87 cell signal intensity at (**A**) day 3 and (**C**) day 6 relative to day 0. CAR-T signal at (**B**) day 3 and (**D**) day 6. Data shown in are means ± s.e.m. *P* values in (**C**) were determined by comparing the area under the curve with one-way ANOVA with Tukey's multiple comparisons test and are <0.0001 (n = 5 for no CAR-T, 13BBζ and CAR 4, df = 12).

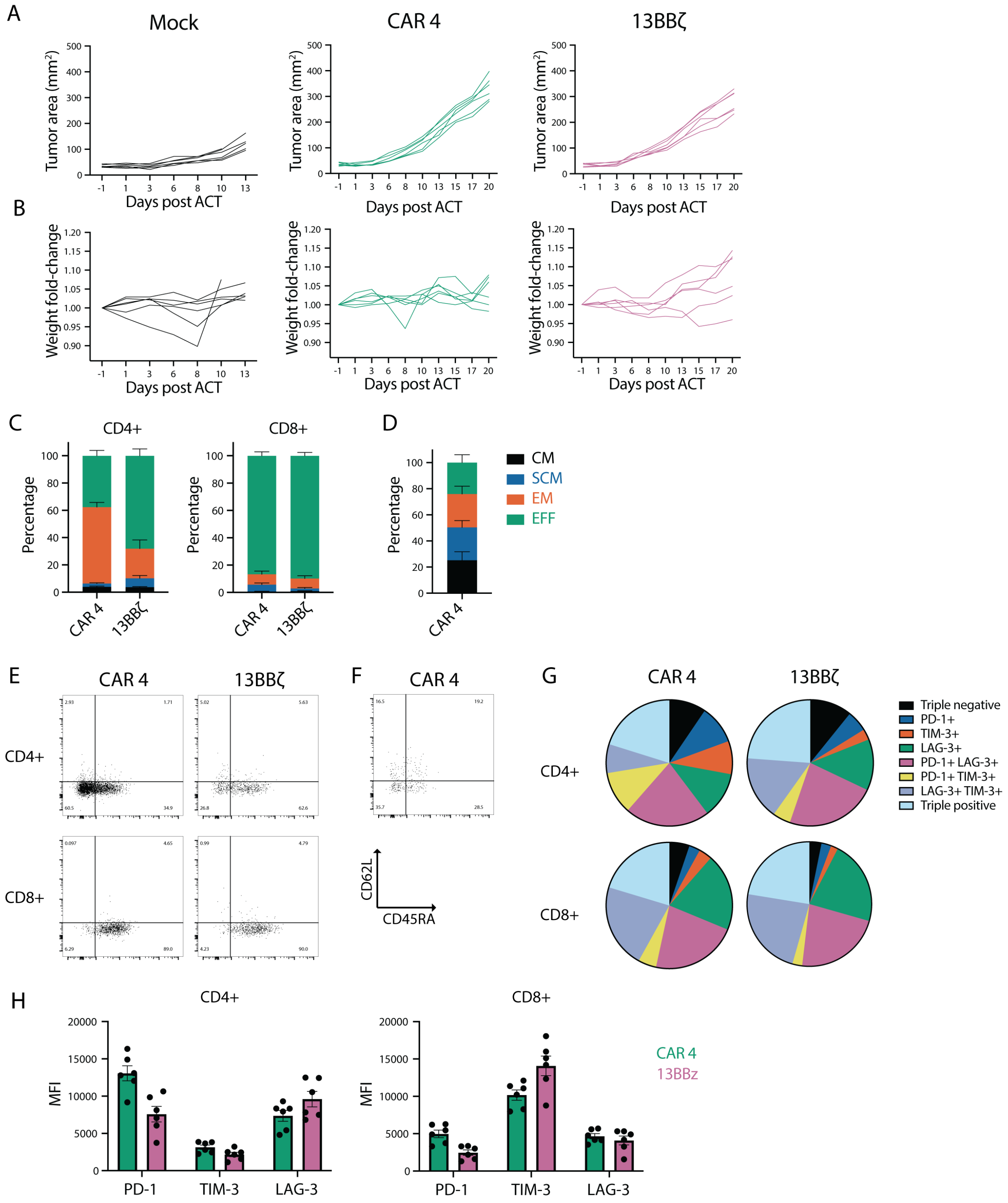


**Supplemental Figure 9.** Related to Figure 6 (donor 3). (**A**) Tumor area and (**B**) weight change following ACT. Data shown are individual values. Distribution of memory phenotype of CD4$^+$ CAR T cells harvested from the (**C**) tumor and (**D**) spleen. FACS plots showing memory phenotype of (**E**) tumor infiltrating and (**F**) splenic CD4$^+$ and CD8$^+$ CAR T cells. (**G-H**) Exhaustion marker expression of tumor infiltrating CD4$^+$ and CD8$^+$ human primary T cells. *P* values in (**H**) were determined using two-way ANOVA with Dunnett's multiple comparisons test.

For PD-1 expression in CD4$^{+}$ T cells, the *P* value is 0.0001 for CAR 4 vs. 13BBζ PD-1 expression. In CD8$^{+}$ T cells, the *P* values are 0.0017 for CAR 4 vs. 13BBζ for CAR 4 vs. 13BBζ TIM-3 expression (n = 6 for CAR 4, and 13BBζ, df = 30).

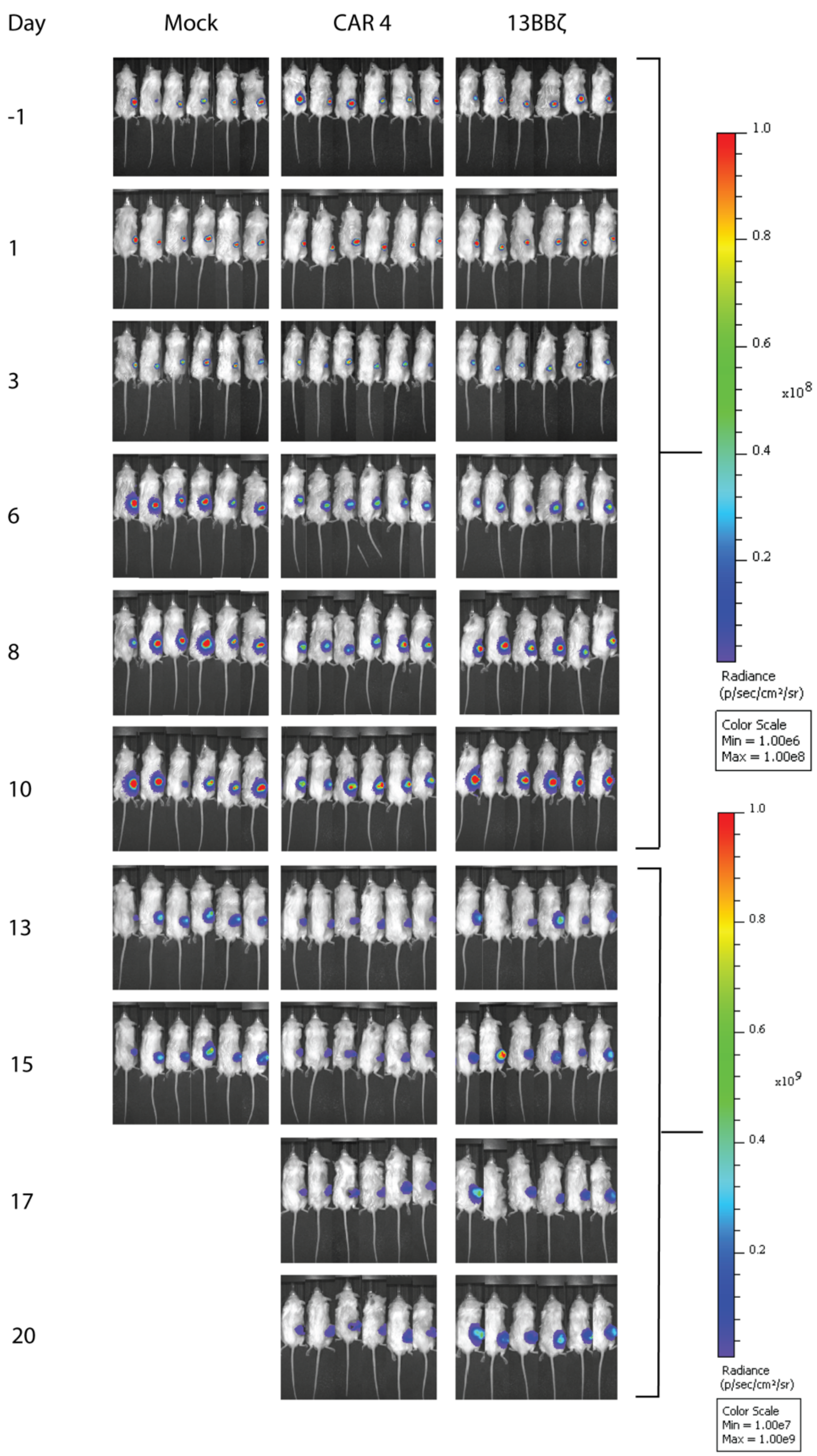


**Supplemental Figure 10**. Related to Figure 7 and Supplemental Figure 9 (donor 3). Luciferase signal from IL13Rα2$^{+}$ U87 cells in mock or CAR-treated mice using IVIS Spectrum imaging after intraperitoneal administration of 0.15 mg of luciferin substrate per gram of body weight.

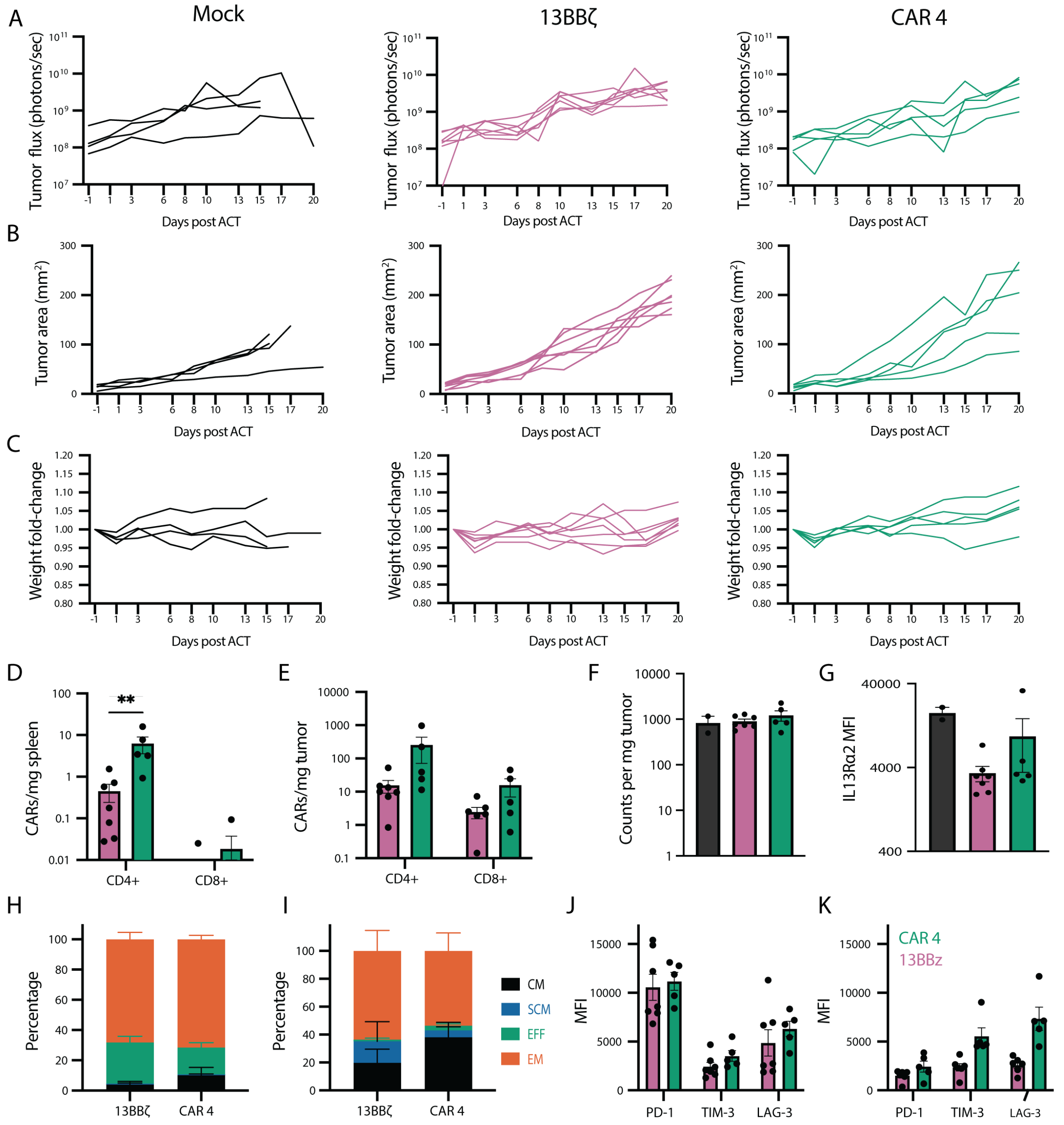


**Supplemental Figure 11.** Related to Figure 7 (donor 4). (**A**) Tumor flux of Flag$^+$ IL13Rα2$^+$ FLuc$^+$ U87 cells implanted subcutaneously in NSG mice. (**B**) Tumor area and (**C**) weight change following ACT. Data shown are for individual mice (n = 5 for CAR 4 and n= 7 for 13BBζ ).(**D**) CD4$^+$ and CD8$^+$ CAR abundance in the spleen and (**E**) tumor. (**F**) Antigen positive tumor cell abundances and (**G**) expression levels. Data shown in (**D-K**) are means ± s.e.m. *P* values in (**D**) and (**E**) were determined using two-way ANOVA with Sidak's multiple comparisons test and are 0.03 and for CAR 4 vs. 13BBζ on day 20 (n = 5 for CAR 4 and n=7 for 13BBζ , df = 18). Mock data in (**F-G**) were dosed with mock transduced donor 3 (n = 1) and donor 4 (n = 1) T cells on the same schedule. Distribution of memory phenotype of CD4$^+$ CAR T cells harvested from the (**H**)

tumor and (**I**) spleen. Exhaustion marker expression of tumor infiltrating (**J**) $CD4^+$ and (**K**) $CD8^+$ human primary T cells.

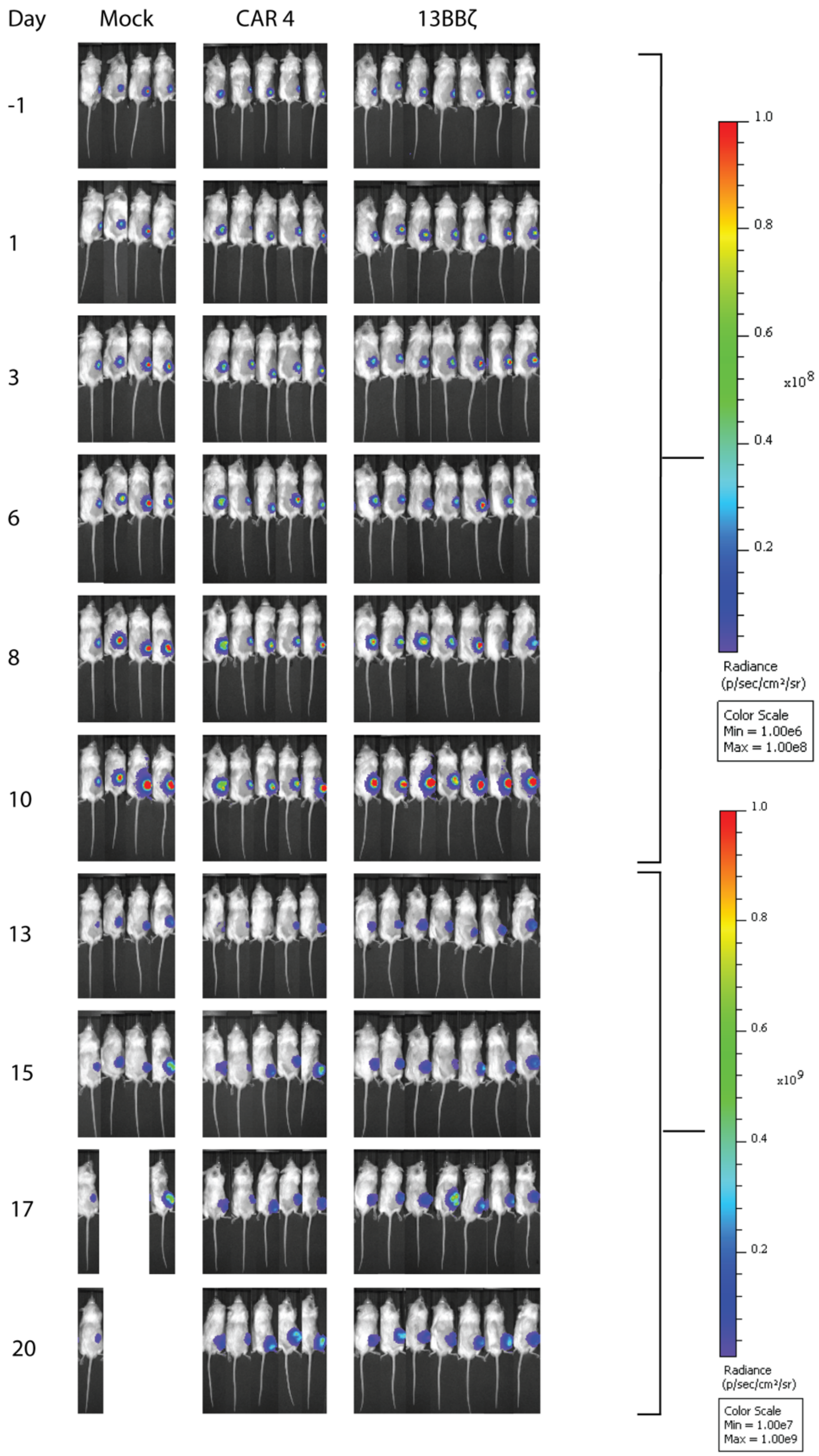


**Supplemental Figure 12**. Related to Figure 7 and Supplemental Figure 11 (donor 4). Luciferase signal from IL13Rα2$^{+}$ U87 cells in mock or CAR-treated mice using IVIS Spectrum imaging after intraperitoneal administration of 0.15 mg of luciferin substrate per gram of body weight.

| **Supplemental Table 1.** Additional amino acid compositions of intracellular signaling domains incorporated into IL-13Rα2-targeted CAR library | | | | | |
|---|---|---|---|---|---|
| ICD | UniProt accession number | Amino acids | ICD | UniProt accession number | Amino acids |
| HHV8 K1 | Q9WHC6 | 248-284 | IL2Rb | P14784 | 266-336,410-427 |
| HHV8 K1 ITAM | Q9WHC6 | 263-280 | IL4Ra | P24394 | 257-321,569-584, 597-612,625-640 |
| EBV LMP2A ITAM | P13285 | 71-88 | *IL7Ra | P16871 | 443-459 |
| EBV LMP2A | P13285 | 471-497 | IL9Ra | Q01113 | 401-416 |
| BLV gp30 | P22507 | 471-515 | *IL12Rb2 | Q99665 | 644-721,792-809 |
| BLV gp30 ITAM | P22507 | 484-501 | *IL21Ra | Q9HBE5 | 513-528 |
| MMTV env ITAM | P03374 | 410-433 | *IL22R1 | Q8N6P7 | 296-309,320-335, 358-399 |
| RRV R1 | Q53D68 | 253-425 | IL23R | Q5VWK5 | 377-445, (478-493,605-620)x2 |
| RRV R1 ITAM 1 | Q53D68 | 396-412 | IFNGR1 | P15260 | 271-320,451-467 |
| RRV R1 ITAM 2 | Q53D68 | 407-424 | EpoR | P19235 | 274-241,360-277, 379-495 |
| ASHV VP7 ITAM | P36325 | 308-324 | IFNAR1 | P17181 | 458-444 |
| | | | IFNAR2 | P48551 | 265-313,329-348, 504-515 |

*STAT domains preceded by EpoR Box1 and Box2 motifs (aa 271-241)
